# GTRspmix: Capturing Heterogeneity of Exchangeabilities Across Sites to Improve Protein Phylogenetics

**DOI:** 10.64898/2026.06.18.729217

**Authors:** Ryo Harada, Edward Susko, Thomas K.F. Wong, Hector Baños, Nhan Ly-Trong, Robert Lanfear, Douglas L. Theobald, Bui Quang Minh, Andrew J. Roger

## Abstract

Site rate and profile mixture models capture the heterogeneity of the amino acid substitution process across sites. However, these models typically use a single matrix of amino acid exchangeabilities and ignore potential heterogeneities of these exchangeabilities across sites. Simply combining multiple exchangeability matrices with rate and profile mixtures leads to a combinatorial explosion of mixture components and a prohibitive increase in free parameters. Here, we introduce GTRspmix, a novel framework that incorporates multiple exchangeability matrices into profile and site rate mixture models while effectively managing model complexity. GTRspmix employs a clustering-based strategy that groups profiles and assigns a distinct exchangeability matrix to each profile cluster. Evaluations using both empirical and simulated datasets demonstrate that GTRspmix fits empirical data significantly better than conventional models, and that overparameterization does not present a problem for sufficiently large alignments. Based on these results, we estimated general-purpose empirical models (SXXpfamCYY series available in IQ-TREE3) from the Pfam database. These general-purpose models not only fit data much better, but they also influence branch length and tree topology estimates, effectively mitigating long-branch attraction artifacts. Because the total number of rate matrices remains manageable, the computational efficiency of the inference is identical to that of conventional profile mixture models (e.g., LG+C60+G4). GTRspmix provides a more realistic and flexible model of protein evolution, offering a robust foundation for the inference of reliable phylogenetic trees.

## Introduction

Amino acid substitution models are of key importance to probabilistic phylogenetic analyses of protein sequences. In these models, the substitution process is modeled as a site-independent Markov process, most commonly assumed to be time-reversible. Although the process was traditionally treated as homogeneous over all sites, this is not biologically realistic; different sites evolve with distinct dynamics depending on their structural or functional roles in the protein. To accommodate this, a number of mixture models have been implemented that account for various kinds of heterogeneity across sites.

The most well-known of these heterogeneities is the differences in evolutionary rate across sites. Rate heterogeneity across sites (RHAS) is typically modeled using a Gamma distribution mixture model, with the shape of the distribution governed by a single parameter, *α* (Yang 1994). The distribution is discretized into several categories of equal probability, with the mean rate of each category used for likelihood calculations. The overall likelihood is thus obtained as the average of the likelihoods for each category. Alternatively, a distribution-free model known as the FreeRate model has been introduced, in which both the rate and weight of each category are estimated from the alignment (Soubrier et al. 2012; Kalyaanamoorthy et al. 2017). This approach often provides a better fit to empirical datasets than the Gamma model.

Another important form of heterogeneity is in the ‘preferred’ amino acids at sites (i.e., preferred amino acid ‘profiles’) that arise from differences in site-specific structural or functional constraints. This amino acid preference heterogeneity across sites (PHAS) has been modeled by mixture models featuring multiple profile classes each of which consists of a vector of 20 equilibrium amino acid frequencies. Lartillot and colleagues implemented a profile mixture model known as the CAT model in a Bayesian framework (Lartillot and Philippe 2004). The CAT model does not fix the number of profiles in advance; instead, it uses a Dirichlet process prior. In maximum-likelihood (ML) analyses, profiles are usually fixed during tree searching to reduce the computational cost. Most commonly, empirical profile sets are used to model PHAS. These include the CXX series (Si Quang et al. 2008) and the UDM series (Schrempf et al. 2020) estimated from databases of alignments. Approaches that estimate profile sets directly from alignments using a composite likelihood were introduced by Susko and colleagues and outperformed empirical profile sets such as CXX (Susko et al. 2018; Williamson et al. 2025). Overall, profile mixture models are known to fit better, and are more robust to phylogenetic artifacts than profile homogeneous models, particularly for deep phylogenetic analyses (Wang et al. 2008; Si Quang et al. 2008; Schrempf et al. 2020; Baños et al. 2023).

There is also potential for heterogeneity in the amino acid exchange rates, known as exchangeabilities, across sites. Different sites may favor different types of amino acid substitutions, and several models have introduced multiple exchangeability matrices to capture this effect. For example, LG4M/LG4X employs different matrices for each rate category (Le et al. 2012), and the EX/EHO/UL series (Le et al. 2008; Le and Gascuel 2010) were designed to account for differences in secondary structure and exposure level between sites. These multi-matrix models generally provide a better fit to empirical data than single-matrix models such as LG+G4 (Le and Gascuel 2008).

While the latter models can be combined with RHAS models (e.g., the Gamma model), they are rarely combined with profile mixture models, mainly due to computational limitations. If multiple exchangeability matrices are simply combined with both rate and profile classes, then every combination of exchangeability matrix, profile, and rate is considered, and the number of mixture components becomes very large. For example, using ten exchangeability matrices in combination with a C60+G4 model results in 10 × 60× 4 = 2400 mixture components. To calculate the site likelihoods in this case, conditional site likelihoods for each site pattern must be separately calculated by the pruning algorithm (Felsenstein 1973) for each of the 2,400 components. Fitting such a model to a large alignment is therefore computationally infeasible both in terms of runtime and memory usage.

An alternative approach would be to employ a distinct exchangeability matrix for each profile. In this case, the total number of components remains the same as in a profile mixture model with a single matrix. Indeed, for nucleotide data, mixture models with multiple exchangeability matrices and frequencies have shown good performance and are recently implemented in IQ-TREE as MixtureFinder (Ren et al. 2025). However, the exchangeability matrix for amino acid data involves 189 free parameters, which is far greater than the five parameters in the nucleotide matrix. Consequently, as discussed by Baños et al. (2024), estimating a unique exchangeability matrix for each of the widely used C60 profiles (i.e., “unlinked” model, following the terminology of Wong et al. (2024)) would require 189× 60 = 11340 parameters, greatly increasing the risk of overfitting compared to nucleotide data. For these reasons, in amino acid sequence analyses, the practical use of both multiple exchangeability matrices and multiple profiles requires strategies to reduce the total number of mixture components as well as the number of exchangeability matrices themselves.

In this study, we propose GTRspmix, a novel framework that incorporates exchangeability heterogeneity across sites (EHAS) into profile and rate mixture models while effectively managing model complexity. To avoid the combinatorial explosion of parameters, we introduce a clustering strategy which groups profiles and assigns a distinct exchangeability matrix to each cluster, rather than to every individual profile or rate category. This reduced set of exchangeabilities can then be optimized based on the data. We assess the performance of GTRspmix using both empirical and simulated datasets, demonstrating that our model fits empirical data significantly better than conventional models and that over-parameterization is not a concern for sufficiently large alignments. Based on this framework, we estimated general-purpose empirical models (SXXpfamCYY series) from the Pfam database. Finally, we evaluated the performance of these new general-purpose models by examining their impact on tree estimation, thereby offering a practical and robust framework for protein phylogenetic inferences. These models have been incorporated into IQ-TREE v3.1.3.

## Methods

### Conventional and Profile Mixture models

The general time-reversible model (GTR) of amino acid substitutions is a Markov process (Tavaré 1986). This process is represented in a 20-by-20 rate matrix *Q* that describes the instantaneous rates of change between amino acid states. Because the process is reversible, the *Q* matrix can be parameterized as the product of a non-negative symmetric exchangeability matrix *S* (i.e., *s*_*ij*_ = *s*_*ji*_) and an amino acid frequency profile *π* = (*π*_1_, …, *π*_20_), with 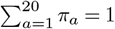:

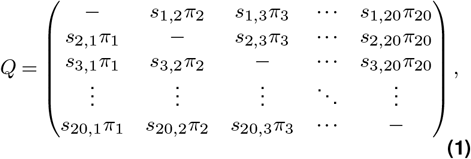

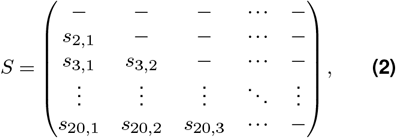

where the diagonal entries of *Q* are defined as 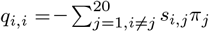 so that the sum of the row is 0. The *Q* matrix is normalized as 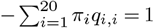 so that the total expected number of substitutions is 1. As a result, there are 189 free parameters per exchangeability matrix, with one entry fixed.

Commonly used profile and rate mixture models such as LG+C60+G4 combine each profile *π*^(*m*)^ and rate category *r*_*k*_ with a single exchangeability matrix *S* to generate multiple rate matrices *Q*. Given an alignment *X* with *N* sites and a tree *T* including branch lengths, the log-likelihood is calculated by following function:

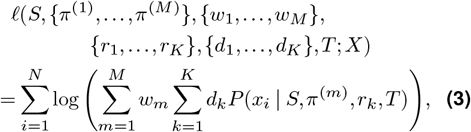

where *w*_*m*_ is the weight of the profile *π*^(*m*)^, *M* is the number of profile mixture components, *d*_*k*_ is the weight of the rate category *r*_*k*_, and *K* is the number of rate heterogeneity categories.

### Motivation for the GTRspmix framework

We hypothesized that sites belonging to similar profiles would evolve under similar exchangeability matrices and, consequently, should be grouped into the same cluster. To justify the clustering of amino acid profiles within our new modeling framework, we conducted a preliminary analysis to assess whether similar profiles correspond to similar exchangeability patterns based on the mutation-selection (MutSel), a mechanistic model of molecular evolution (Kimura 1962). According to the MutSel model, the profile and exchangeability matrix are derived from the relative fitness of each amino acid and the profile and exchangeability matrix without selection pressure (Kimura 1962; Halpern and Bruno 1998; Yang and Nielsen 2008; Rodrigue et al. 2010; Wang et al. 2014). Following the MutSel model, we randomly generated sets of relative fitnesses to derive 60 pairs of profiles and their corresponding exchangeability matrices. We calculated pairwise distances for both the profiles and the matrices, partitioning them into three categories of magnitude (low, mid, and high) based on tertiles of the distribution. Of the unique pairwise comparisons, 78% exhibited concordance between the profile and exchangeability distance categories (see Table 7). These results support the assumption that sites with similar profiles tend to evolve under similar exchange-ability matrices. The detailed derivation of the MutSel model and description of these analyses is provided in the Appendix.

In this study, we introduce the GTRspmix model, which uses multiple exchangeability matrices together with profile and rate heterogeneity across sites. The profiles are divided into *C* clusters, and each cluster *c* has its own exchangeability matrix *S*_*c*_. The log-likelihood is then calculated as follows:

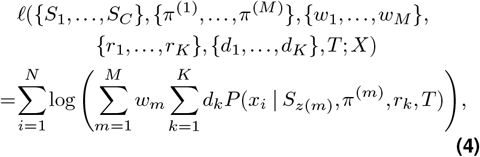

where *z*(*m*) represents a function that returns the cluster label *c* ∈ *{*1,…, *C}* to which the profile *π*^(*m*)^ belongs.

### Model Estimation Methodology

#### Profile clustering based on site posterior probability co-occurrences

To estimate the parameters of the GTRspmix model, profiles must first be clustered. Let *H* = {*π*^(*H*,1)^, …, *π*^(*H,M*)^} denote the set of profiles to be clustered. One way to extract the *C* clusters of profiles, each sharing the same matrix *S*_*c*_, is to cluster the profiles based on the similarity of 20 amino acid frequencies using `k-means++` (Arthur and Vassilvitskii 2006) or other clustering methods. In addition to this similarity-based clustering, we propose another approach based on the posterior probabilities of profile classes given each site for two different profile sets.

The posterior probability for component *m* of a profile mixture model given data at site *i* is computed as

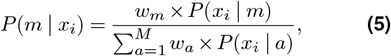

where the *w*_*m*_ is the weight of the mixture component *m* and *P* (*x*_*i*_ |*m*) is the probability of site pattern *x*_*i*_ given component *m*.

We propose a novel clustering approach, Site Posterior Probability Co-occurrence (SPPC), which groups a larger set of profiles (denoted as *H*) into at most *C* clusters based on a smaller set of profiles (*J* = {*π*^(*J*,1)^, …, *π*^(*J,C*)^}, where *C* < *M*). SPPC has following steps:

1. For a given alignment, we independently calculate the posterior probabilities of each mixture component (profile) for each site under both profile sets *H* and *J*. Each site is then assigned to the profile that exhibits the maximum posterior probability in each set.
2. For each profile in the larger set *H*, we restrict attention to all sites assigned to the profile.

Among these sites, we find the most frequently co-occurring profile (i.e., the mode) from the smaller set *J*. This provides a many-to-one mapping from *H* to *J*. Finally, profiles in *H* that are mapped to the same profile in *J* are grouped together.

To clarify the SPPC algorithm, we provide a simple example using a toy alignment (Fig. 1). Suppose that we have a larger profile set *H* with five profiles (*π*^(*H*,1)^ to *π*^(*H*,5)^) and a smaller reference set *J* with three profiles (*π*^(*J*,1)^ to *π*^(*J*,3)^). For an alignment consisting of 10 sites, each site is first assigned to its best-fitting profile in both sets based on the maximum posterior probability.

**Figure 1.**
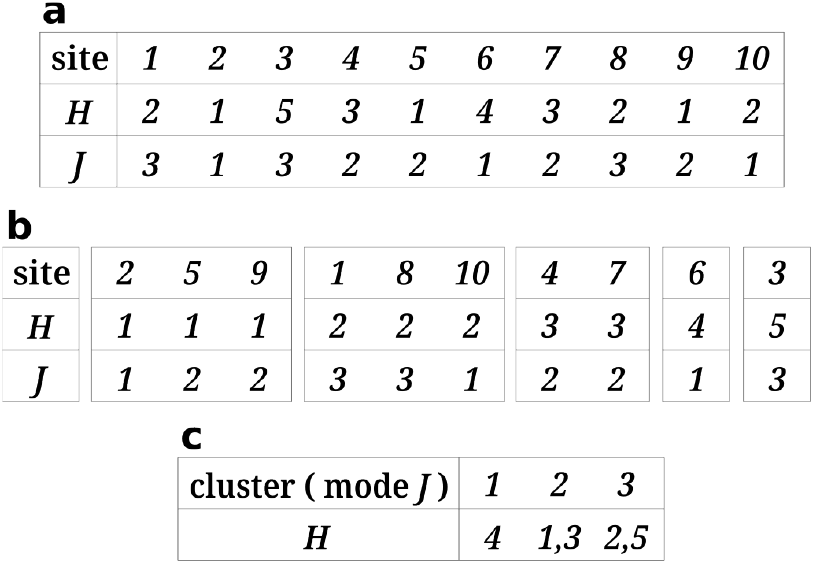
An example of the Site Posterior Probability Co-occurrence (SPPC) clustering algorithm. A larger profile set *H* is clustered using a smaller profile set *J* over a 10-site alignment. a) Each site is assigned to a specific profile in *H* and *J* based on the maximum posterior probabilities. The numbers in the rows for *H* and *J* represent the profile indices. b) Sites assigned to each profile in *H* are aggregated to identify the co-occurring profiles in *J*. c) Profiles in *H* are grouped into clusters based on the most frequent *J* profile, where “mode *J*” indicates the most frequently co-occurring *J* profile associated with each cluster.

For instance, among the sites assigned to *π*^(*H*,1)^, the profile *π*^(*J*,2)^ co-occurs most frequently (two times), while *π*^(*J*,1)^ occurs only once; thus, *π*^(*H*,1)^ is mapped to *π*^(*J*,2)^. Following the same majority rule, *π*^(*H*,2)^ and *π*^(*H*,5)^ are mapped to *π*^(*J*,3)^, *π*^(*H*,3)^ is mapped to *π*^(*J*,2)^, and *π*^(*H*,4)^ is mapped to *π*^(*J*,1)^. As a result, the five profiles in *H* are naturally partitioned into three distinct clusters corresponding to the profiles in *J*: *{π*^(*H*,4)^*}* associated with *π*^(*J*,1)^, *{π*^(*H*,1)^, *π*^(*H*,3)^*}* associated with *π*^(*J*,2)^, and *{π*^(*H*,2)^, *π*^(*H*,5)^*}* associated with *π*^(*J*,3)^.

#### Approximating the Expectation Maximization (EM) Algorithm

IQ-TREE (Minh et al. 2020; Wong et al. 2026) provides a way to obtain a single exchangeability matrix by ML estimation, but it cannot directly optimize multiple exchangeability matrices in our mixture model. To overcome this limitation, we adopt a modified version of the Expectation-Maximization (EM) algorithm (Dempster et al. 1977). Specifically, our approach optimizes a single exchangeability matrix for each cluster of profiles by maximizing the expected log-likelihood of the corresponding component.

Under the assumptions of the GTRspmix model, each site belongs to one of the exchangeability matrices, and each matrix is associated with a cluster of profiles. We introduce *y*_1_, …, *y*_*N*_ as the unobserved cluster assignment labels, which map each site to a single cluster *c*. The log-likelihood of the complete data is

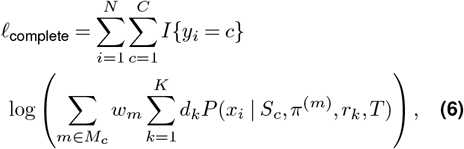

where *M*_*c*_ is the set of profile labels belonging to cluster *c* and *I* {*y*_*i*_ = *c*} is an indicator function that returns 1 when the class label *y*_*i*_ of site *i* is equal to the cluster index *c*, otherwise 0. Because we do not know the class labels, the log-likelihood that we actually want to maximize is Eq. (4).

The EM algorithm provides a systematic way to address this difficulty. It iteratively constructs an objective function based on the expected complete-data log-likelihood and updates parameters to maximize the expectation. The expected log-likelihood is calculated as

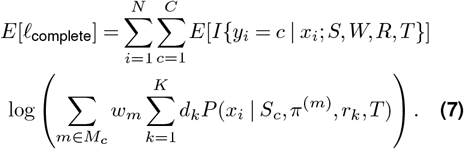

Since, the expectation, *E*[*I*{*y*_*i*_ = *c*| *x*_*i*_; *S, W, R, T}*], is taken with respect to the distribution of latent labels *y*_*i*_, this term can be replaced by the posterior probability of cluster *c* given site data *x*_*i*_.

The GTRspmix model estimation employing the EM approximation consists of the following four steps, which are detailed below (Fig. 2).

**Figure 2.**
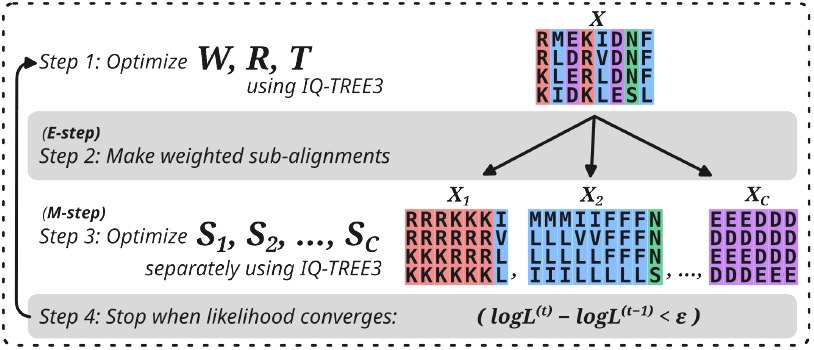
Workflow of optimization of GTRspmix using EM approximation. Here, *W* denotes the profile weights, *R* represents the rates and weights for rate heterogeneity, *T* indicates branch lengths, and *S* is a set of exchangeability matrices.

#### Step 1: Initialization and estimation of the other parameters

First, given current exchangeability matrices 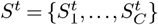, optimize the current parameters, *T* ^*t*^, *W* ^*t*^, and *R*^*t*^, using Eq. (4). Here, *W* denotes the profile weights, *R* represents the rates and weights associated with rate heterogeneity, and *t* indicates the iteration. This is the usual model estimation with fixed exchangeability matrices and profiles using IQ-TREE. Note that the initial parameters for all exchangeability matrices 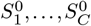 are the same single user-defined matrix, so the first iteration does not differ from Eq. (3).

#### Step 2: The E step

Using the current parameters, the posterior probability for cluster *c* given the data *x*_*i*_ at site *i* is computed as

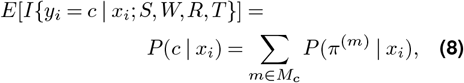

where *P* (*c*| *x*_*i*_) is the sum of the posterior probabilities for profiles belonging to cluster *c*. This step is performed by summing the IQ-TREE output computed with the “`-wspm`” option in the same run as step 1.

#### Step 3: Approximation of the M step

IQ-TREE does not directly allow maximization of the expected complete log-likelihood but does allow ML estimation of a single exchangeability matrix for an alignment. We approximate the M step with ML estimation for an alignment. We introduce a scaling factor *b* (e.g. 10, 10^2^, 10^3^) and define the count *N*_*c*_(*x*_*i*_) for each cluster *c* as

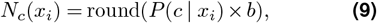

where `round()` denotes the standard rounding function to the nearest integer.

The expected log-likelihood for each exchangeability matrix component *c* can be approximated by replacing the fraction weights *P* (*c* | *x*_*i*_) with the scaled integer:

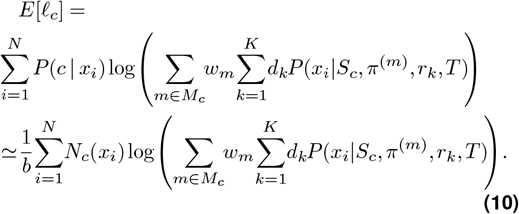

For each *c*, we obtain new exchangeability parameters 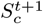 that maximize *E*[*l*_*c*_] given the current parameters. Technically, we generate a sub-alignment *X*_*c*_ for each cluster *c* using “soft-partitioning.” The site data *x*_*i*_ is then allocated *N*_*c*_(*x*_*i*_) times to the sub-alignment of cluster *c*. Thus, the sub-alignment *X*_*c*_ constructed in this way provides a discrete approximation of the contribution of each site to the expected log-likelihood. We optimize a single exchangeability matrix with profiles in the corresponding cluster for each of the weighted sub-alignments using GTRpmix implemented in IQ-TREE (Baños et al. 2024). The scaling factor *b* determines how many decimal places of the posterior probabilities are effectively represented; increasing *b* improves precision. As *b* becomes sufficiently large, the approximated log-likelihoods converge to the actual values.

#### Step 4: Convergence check

Update the exchange-ability matrices with the newly obtained 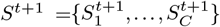 If *l*(*S*^*t+1*^) − *l*(*S*^*t*^) < *ϵ* (where *ϵ* is a user-defined convergence threshold), the model optimization finishes and reports final *S*. Otherwise, go to Step 1 with the new matrices

It is worth noting that our EM approximation is similar to the approach used in estimating the LG matrix and the LG4M/LG4X models (Le and Gascuel 2008; Le et al. 2012), which constructed a sub-alignment for each rate category based on the maximum posterior probability assignment. However, their method differs in that sites are fully partitioned into sub-alignments according to the maximum posterior probability, whereas in our method, each site contributes to all mixture components, with the allocation determined by scaled and rounded posterior probabilities. By allowing a larger scaling factor, our routine provides a more accurate discrete approximation of the EM algorithm, thereby ensuring that it effectively maximizes the likelihood for the observed data. However, larger values of *b* entail higher computational costs in terms of storage and optimization time. As a practical heuristic to balance accuracy and efficiency, we fixed *b* = 10 for all model estimations throughout this study. This choice accounts for posterior probabilities down to the first decimal place while effectively omitting sites with negligible support (less than 0.05), thereby saving computation time.

### Model Estimation and Evaluation

After implementing the GTRspmix script, we evaluated the performance of the GTRspmix model both with empirical and simulation datasets. The properties of the empirical datasets used in this study are shown in Table 1.

**Table 1.** Empirical Datasets used in this study.

| Dataset | Proteins | Sites | Taxa | References |
| --- | --- | --- | --- | --- |
| Eukaryota | 240 | 76,840 | 78 | Baños et al. (2024) |
| Pfam train | 742 | 231,363 | 130,684 | El-Gebali et al. (2018) |
| Pfam test | 742 | 245,649 | 124,881 | El-Gebali et al. (2018) |
| Microsporidia | 133 | 24,291 | 40 | Brinkmann et al. (2005) |
| Nematode | 146 | 35,366 | 37 | Lartillot et al. (2007) |
| Platyhelminthes | 146 | 35,367 | 32 | Lartillot et al. (2007) |

#### Holdout test on the Eukaryota dataset

To test GTRsp-mix under different numbers of clusters and clustering schemes, we used the Eukaryota alignment, a dataset comprising 240 protein-coding genes with 78 representative eukaryotic taxa originally compiled from the PhyloFisher database (Tice et al. 2021) and used in the GTRpmix study (Baños et al. 2024). We estimated a tree from this alignment using IQ-TREE under the ELM+C60+G4 model (Si Quang et al. 2008; Baños et al. 2024) with optimized mixture weights (“`-mwopt`”), where the ELM matrix was optimized specifically for this alignment. This fixed ML tree was used for the estimation and evaluation of the following GTRspmix models.

Based on the Eukaryota alignment and the ML tree, we estimated sets of custom profiles with varying numbers of classes (2, 4, 6, 8, 10, 20, 30, 40, 50, and 60) using a MEOW pre-released version with the “`-p IQ`” option (Williamson et al. 2025). The MEOW is an extension of the MaMMAL approach (Susko et al. 2018), that makes use of all sites and various partitioning schemes to improve profile estimation. For each MEOW model, we calculated posterior probabilities for each profile at every site using the ELM matrix and the G4 RHAS model with “`-wspm`” flag in IQ-TREE (v2.3.4), and clustered MEOW60 profiles based on the SPPC approach with each of the other MEOW models. Then the MEOW60 profiles were clustered into different numbers of groups; the number of resulting clusters in each case corresponded to the number of exchange-ability matrices in the respective GTRspmix model (i.e., each cluster was assigned a single exchangeability matrix). For MEOW40 and MEOW50, three profiles did not co-occur with any profiles in MEOW60; the resulting MEOW60 profiles were clustered into 37 and 47 groups, respectively. As an alternative approach, we also clustered MEOW60 profiles using the `k-means++` algorithm (Arthur and Vassilvitskii 2006) with a range of K values.

The Eukaryota alignment was randomly split into two sub-alignments of 38,420 sites each; one for training and the other for testing. GTRspmix models were estimated using the training alignment and the various MEOW60 clustering results, and their performance was subsequently evaluated on the test alignment. All parameters, including branch lengths, Gamma shape parameter *α*, and mixture weights are estimated using the training alignment and fixed in evaluation on the test alignment. As conventional models, we included the POISSON matrix, which has equal exchangeabilities between all amino acid states, and the GTRpmix matrix, in which a single exchangeability matrix is linked to all profiles. All models used MEOW60 profiles and the G4 RHAS model. Here, we denote our GTRspmix models as SXXMYY+G4, where XX indicates the number of exchangeability matrices (i.e., clusters), and YY denotes the number of MEOW profiles.

We evaluated model performance primarily by comparing log-likelihoods on the test alignment. This approach directly assesses the model’s ability to generalize to unseen data, as over-fitted models will naturally yield poorer test likelihoods. For conventional compar-isons, we also calculated the Akaike Information Criterion (AIC; Akaike 1974) and Bayesian Information Criterion (BIC; Schwarz 1978) for the training alignments. To assess whether alternative models showed significantly different fits relative to the best-fitting model on the test data, we compared their site-wise log-likelihoods. For a given site *i*, let *ℓ*_1*i*_ and *ℓ*_2*i*_ denote the log-likelihoods under the best-fitting model (model 1) and an alternative model (model 2), respectively. We defined the sitewise difference as *d*_*i*_ = *ℓ*_1*i*_− *ℓ*_2*i*_. We then performed a paired Z-test to test the null hypothesis that the mean difference was 0.

#### Assessing overparameterization using simulated data

To assess the potential overparameterization of the GTRspmix model when an excessive number of exchange-ability matrices is specified, we conducted simulation analyses. Training and test alignments, each containing 10,000 sites, were simulated under the LG+C10+G4 model with uniform C10 mixture weights (each profile contributing 10%) and a Gamma shape parameter *α* of 1.0 using AliSim in IQ-TREE 3.0.1 (Ly-Trong et al. 2022; Wong et al. 2026). The 32-taxon tree (Supplementary Fig. S1) was used for data generation, model training, and model evaluation. The training alignment was used to estimate the S01C10+G4 (equivalent to GTRpmix+C10+G4), S02C10+G4, S03C10+G4, S05C10+G4, and S10C10+G4. In this analysis, the number following S denotes the number of exchange-ability matrices, and all models were combined with C10 profiles. The S10C10+G4 model has one exchange-ability matrix for each profile, and is equivalent to the IQ-TREE command “`-m GTR20+C10+G4`” where the “20” in GTR20 denotes the number of amino acid states. For optimization of GTRspmix models, the C10 profiles were clustered using `k-means++`.

To examine the effect of alignment length, we repeated these analyses with downsampled training alignments of 100, 200, 500, 1,000, 2,000, and 5,000 sites randomly selected from the original training alignment. For performance comparison, we also evaluated the other models: (i) the true model (LG+C10+G4), (ii) alternative exchangeability matrices (Qmaker series, WAG, JTT, and POISSON) with C10+G4 (Jones et al. 1992; Whelan and Goldman 2001; Minh et al. 2021), and (iii) the single-profile LG+F+G4 model. In all evaluations, all parameters were estimated from training alignments and fixed in evaluations on the test alignment.

#### Optimization and comparison of general-purpose models using the Pfam database

Following the evaluation on the Eukaryota alignment, we optimized general-purpose GTRspmix models using the Pfam database version 31 (El-Gebali et al. 2018). We downloaded the Pfam alignments used for Q.pfam estimation, re-aligned them using MAFFT L-INS-i model (Katoh and Standley 2013), and trimmed them using trimAl with “`-gt 0.5`” option (Capella-Gutiérrez et al. 2009). From these, we selected 1,484 alignments that contained more than 50 taxa and 200 sites for downstream analyses. For each alignment, we inferred the ML tree using IQ-TREE v3.0.1 under the Q.pfam+C60+G4 model with optimized weights, and used these ML trees for optimization and evaluation of GTRspmix models. The alignments and their corresponding trees were split into two sets of 742 alignments each, one for training and the other for testing.

Due to computational limitations, we focused on optimizing GTRspmix models using C60 profiles with 10 and 30 exchangeability matrices determined by SPPC clustering. We chose these numbers of profiles and exchangeability matrices based on their good performance in the evaluation on the Eukaryota alignment (Fig. 3 and Supplementary Table S1). For C60 clustering, we computed site-wise posterior probabilities for the C10, C30, and C60 profile sets. We clustered the C60 profiles into 10 and 30 groups using the SPPC method based on co-occurrence between C60 vs. C10 and C30, respectively (Supplementary Figs. S2–S4). In the case of C30-based clustering, one profile (profile 6 in IQ-TREE ordering) out of 30 did not co-occur with any of the C60 profiles, resulting in 29 clusters. In addition, one cluster consisted solely of profile 21, which had a fairly low weight (2 × 10^−7^), making exchangeability matrix optimization for this cluster impossible. Therefore, we excluded this cluster from model training and optimized 28 exchangeability matrices on top of the remaining 59 profiles (C59). After optimization of 28 exchange-ability matrices on 59 profiles, we also tested an alternative model that added a class combining the POISSON matrix and the profile 21 from C60, which had been removed before optimization. In addition, when processing a large number of alignments or a huge alignment, computational resource limitations can make it impractical to use the C60 models. To address such cases, we also optimized computationally efficient models in which each profile of C10, C20, and C30 was assigned its own exchangeability matrix.

**Figure 3.**
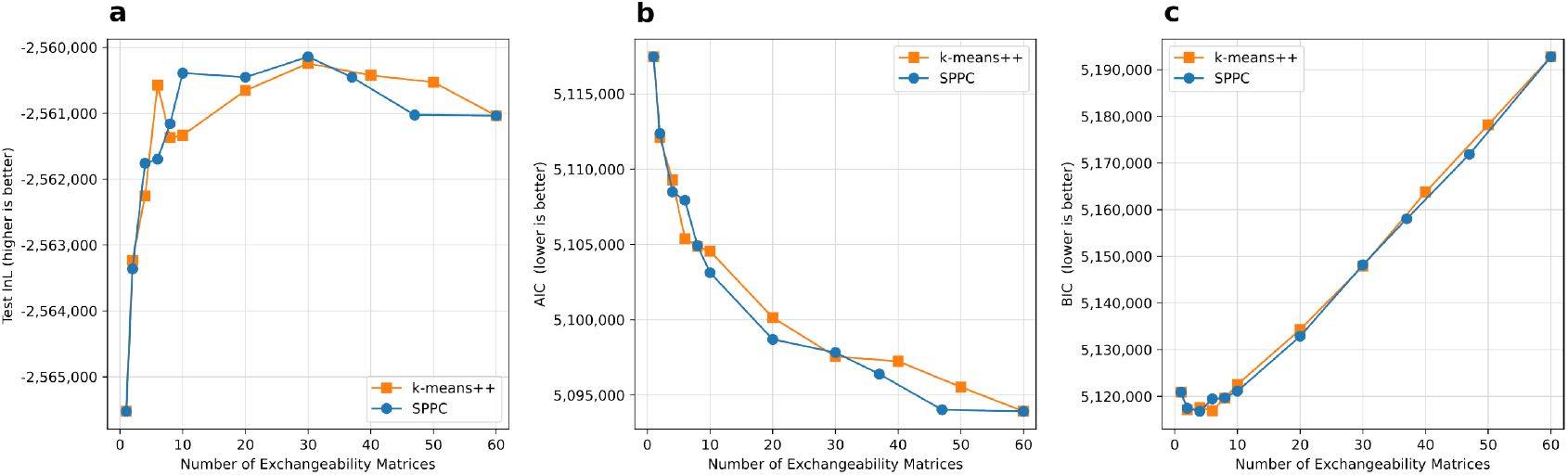
Performance of GTRspmix models on the Eukaryota dataset. The x-axis in all panels represents the number of optimized exchangeability matrices. a) The log-likelihood of the test alignment. b) Akaike Information Criterion (AIC) assessing model fit to the training alignment. c) Bayesian Information Criterion (BIC) assessing model fit to the training alignment. The lines marked with circles and squares represent SPPC and k-means++ clustering, respectively.

As additional benchmarks, we included the Q.pfam+G4 model (a single-profile model), the Q.pfam+C60+G4, and the GTRpmix+C60+G4 model with optimized weights. For the GTRpmix+C60+G4 model, we optimized a single exchangeability matrix with C60 profiles using the Pfam training dataset; we call this GTRpmix-optimized matrix G.pfam. In this study, our naming convention for general-purpose GTRspmix models optimized using the Pfam database is SXXpfam-CYY, based on the number XX of exchangeability matrices and the number YY of profiles (e.g., S10pfamC10, S10pfamC60, and S28pfamC59). All of our model optimizations included the G4 model.

To evaluate the performance of the GTRspmix models and conventional models, we compared their log-likelihoods on the Pfam test dataset. In the Pfam dataset, training and test data were partitioned at the gene level to ensure sufficient alignment length per gene. Consequently, the training and test sets comprised different trees. For model evaluation, branch lengths, the shape parameter *α*, and mixture weights were estimated using the test data. In addition to the log-likelihood comparisons, we also checked branch length estimates. For each pair of models, we calculated the correlation coefficients of the estimated branch lengths on the Pfam test tree set. To assess the impact of GTRspmix models on phylogenetic topology inference, we randomly selected 100 alignments from the Pfam test database and conducted tree searches to obtain the ML trees under each of the newly estimated and conventional models. For each pair of models, we calculated the normalized Robinson-Foulds (nRF) distance between the ML trees of each alignment (Robinson and Foulds 1981). To visualize the relationships between models and their topological estimates, we constructed hierchical clusterings (Unweighted Pair Group Method with Arithmetic Mean; UPGMA; Sokal and Michener 1958) using element-wise median and mean nRF distance matrices derived from the 100 alignments. To evaluate the variability with which models will tend to produce similar topological estimates, we calculated the frequencies with which each cluster in the overall hierarchical clustering occurred across 100 individual hierarchical clusterings over individual gene trees (analogous to bootstrap support).

#### Comparison using empirical LBA datasets

In addition to the evaluation on the Pfam test data, we assessed the performance of general-purpose GTRspmix models on three empirically recognized long-branch attraction (LBA) datasets previously used in Wang et al. (2018), Susko et al. (2018), and Baños et al. (2024). Each dataset is associated with both a “true” reference tree and an “artifactual” LBA tree. Although the absolute true tree is essentially unknown for empirical data, these widely accepted reference trees are hereafter referred to as the “true tree” for simplicity.

1. Microsporidia dataset — Early analyses attempting to place Microsporidia within the eukaryotic tree were artifact-prone because of the extremely divergent gene sequences of these obligate intracellular parasites. It is now known that the true tree, recovered under the LG+C20+F+G4 model, places Microsporidia as sister to Fungi, whereas the LBA tree, reconstructed with the LG+F+G4 model, places them as sister to the archaeal out-group (Brinkmann et al. 2005).
2. Nematode dataset — For this dataset, the true tree, inferred under the LG+C20+F+G4 model, places nematodes as sisters to arthropods, while the topology inferred under the LG+F+G4 model is an LBA artifact (Lartillot et al. 2007).
3. Platyhelminthes dataset — For this dataset, the true tree recovered under the GTR+CAT+G4 model, places platyhelminthes within Protostomia, whereas the LBA tree inferred under the LG+F+G4 model places them within Coelomata (Lartillot et al. 2007).

For each dataset, we fitted all models to both the true tree and the LBA tree, and obtained their log-likelihoods. Each model will show some level of support for or against the true tree. We wanted to assess whether these differences in support might be explained by sampling variation among sites or whether there was significant evidence for a real mean difference over two models in their level of support for the true tree. To do this, we calculated site-wise log-likelihood differences (True− LBA) for the two models, *d*_1*i*_ and *d*_2*i*_, for a given site *i* and models 1 and 2. We then performed a paired Z-test to test the null hypothesis that the mean difference between the two models was 0, treating the difference in support, *d*_*i*_ = *d*_1*i*_ −*d*_2*i*_, as the data for the test.

We further evaluated relative model fit using an approximation of the extant sequence reconstruction (ESR) model comparison framework proposed by Sennett and Theobald (2024) which is a type of cross-validation (CV). In ESR, the probability of the observed amino acid for each site and taxon is calculated conditional upon the amino acids for all other taxa at that site. Large probabilities on average are taken as evidence for a good model, and a number of measures are considered; here, we use the summed log-probabilities. In the site-wise CV version of ESR, for every observed (extant) amino acid position in every sequence, parameters are re-estimated without that amino acid. This makes predictions independent of the amino acid for the taxon and site of interest, but it is computationally intensive. Our approximation uses parameters estimated from the original full alignment. We expect similar results to ESR with this approximation because the deletion of a single amino acid will not usually affect parameter estimates in a substantial way.

For the LBA datasets, we fitted general-purpose models with the true trees and compared them using ESR. As a metric of ESR-based model comparison, we used the predictive log-probabilities of the observed extant se-quences under a given model, defined as

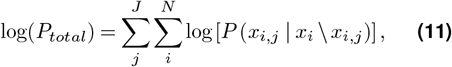

where *J* denotes the number of taxa in the tree, and *N* is the number of alignment sites. The variable *x*_*i,j*_ represents the amino acid observed at site *i* in taxon *j*, and *x*_*i*_ is the vector of amino acids at site *i* across all taxa. The operator “\” denotes the exclusion of *x*_*i,j*_ from the set *x*_*i*_. Thus, *P* (*x*_*i,j*_ |*x*_*i*_ *\x*_*i,j*_) corresponds to the probability that amino acid for site *i* in taxon *j* matches the observed state, calculated conditional upon the amino acids for all other taxa at the site.

## Results

### Using Multiple Exchangeability Matrices Improves Model Performance

To evaluate the performance of GTRspmix models as well as the effects of the number of exchangeability matrices and clustering methods, we conducted a hold-out test using the Eukaryota alignment of 240 eukaryotic genes. The holdout test showed that GTRspmix models, which employ multiple exchangeability matrices, substantially improved the log-likelihoods for both the training and the test datasets compared to conventional models that use a single matrix (Fig. 3 and Supplementary Table S1). For the training dataset, log-likelihood increased as the number of exchangeability matrices (and thus the number of free parameters) increased. The model selection criteria showed different preferences. The AIC (Akaike 1974) favored the model with 60 exchangeability matrices (S60M60+G4; Fig. 3b), which estimated one exchangeability matrix for each of the MEOW60 profiles and was equivalent to the “`GTR20+MEOW60+G4`” model in the IQ-TREE command. In contrast, the BIC (Schwarz 1978), which penalizes model complexity more strongly, preferred the model with four exchangeability matrices (S04M60+G4; Fig. 3c). The log-likelihood of the test dataset was maximized by the model with 30 exchangeability matrices (S30M60+G4), and the model with 10 exchangeability matrices (S10M60+G4) ranked second (Fig. 3a).

Two clustering approaches, SPPC and `k-means++`, showed no major performance differences, and both clustering-based GTRspmix models outperformed single-matrix models. However, for the top-performing model (S30M60+G4) and the second-best model (S10M60+G4), the SPPC approach yielded slightly better test log-likelihoods than `k-means++`. Therefore, we adopted SPPC as the clustering method for subsequent analyses.

Notably, improvements in test log-likelihood seemed to plateau after the number of exchangeability matrices exceeded 10. The difference in test log-likelihood be-tween the best model (S30M60+G4 with SPPC) and the second-best model (S10M60+G4 with SPPC) was only 6.48 × 10^−3^ log-likelihood units per site. We conducted a Z-test using site-wise log-likelihoods from the test alignment, which did not detect significant differences between S30M60+G4 and S10M60+G4 models (*p* = 0.2356). These results indicate that, for the Eukaryota alignment, the S10M60+G4 model is a reasonable alternative to the more parameter-rich S30M60+G4 model when one prioritizes avoiding overparameterization.

However, even for models with numbers of exchange-ability matrices greater than 30, the decrease in test log-likelihood was not major (1.03 × 10^−2^ points per site between 30 and 60 matrices), and performance remained better than those of single-matrix models or models with fewer matrices. This suggests that overparameterization may not be a serious concern for GTRspmix models.

### Overparameterization Is Not a Serious Problem for Large Alignments

To evaluate the risk of overparameterization, we tested an extreme case using simulated data. The training and test alignments were generated under the LG+C10+G4 model, in which EHAS was absent, meaning that all sites shared a single exchangeability matrix (LG). Under this condition, when GTRspmix with multiple exchange-ability matrices is optimized on the training alignment, the model is expected to overfit the training data, resulting in a lower log-likelihood for the test alignment compared with the single matrix models.

As expected, the test log-likelihood decreased as the number of exchangeability matrices increased, particularly for shorter alignments (Fig. 4). The horizontal lines provide references. Each line corresponds to replacing the LG exchangeability matrix in the simulation model with a different empirical one. Since there is only one exchangeability matrix for each line and none of its parameters is estimated, these models have relatively few parameters. It is known that the JTT and WAG matrices are similar to the LG matrix, so performance similar to theirs should be considered satisfactory. When the training alignment length reached 2,000 sites, even the S10C10+G4 model, which has nine more exchange-ability matrices than the true model, performed similarly to the JTT+C10+G4 model, although it had a slightly lower log-likelihood score. For larger training alignments (5,000 and 10,000 sites), S10C10+G4 consistently outperformed JTT+C10+G4. Given the similarity between the JTT and LG matrices, this decline of test log-likelihood was relatively minor. Furthermore, for the 10,000 sites training alignment, S10C10+G4 out-performed all other models except LG+C10+G4 and Q.pfam+C10+G4, the first being the simulation model and the second having exchangeabilities that are very similar to those of the LG matrix (Le and Gascuel 2008; Minh et al. 2021).

**Figure 4.**
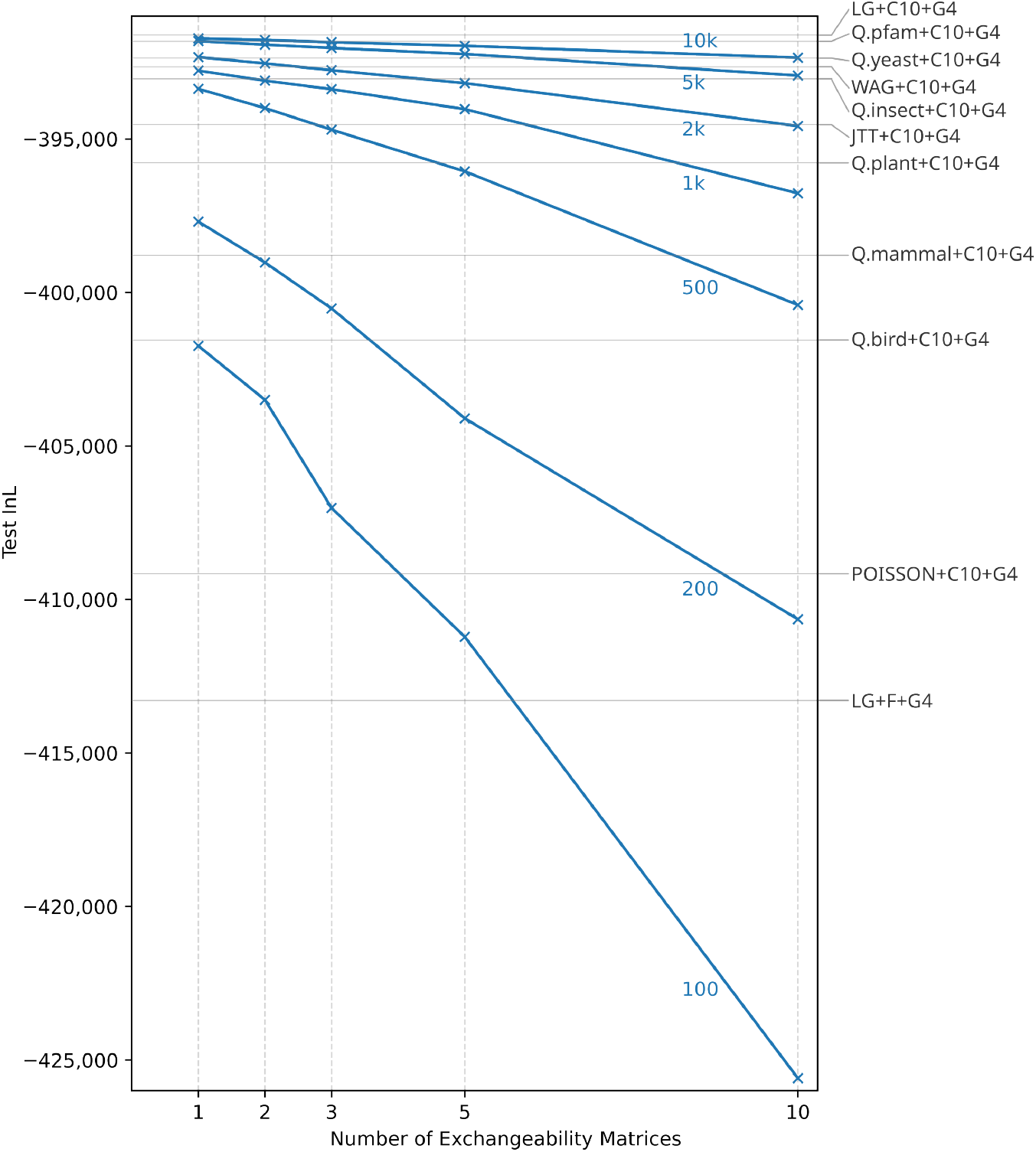
Performance on the test alignment when overparameterization is induced by estimating GTRspmix from sequences generated under the LG+C10+G4 model. The x-axis shows the number of optimized exchangeability matrices, and the y-axis shows the log-likelihood of the test alignment. Results for different training-alignment lengths used for model estimation are shown as lines marked with × symbols, with numbers indicating the corresponding training-alignment lengths. The horizontal lines indicate the test log-likelihoods obtained using each of several empirical models; the corresponding model names are shown on the right side of the figure.

These results indicate that for large datasets, even if the data were generated by a single exchangeability matrix, the fitted GTRspmix models with large numbers of excess parameters do not show signs of overfitting. In the holdout test using the empirical Eukaryota alignment (described above), the test log-likelihood showed an increasing trend (Fig. 3a) rather than the decreasing trend observed in the simulation (Fig. 4). This suggests that the increasing trend in Fig. 3a is not a statistical artifact of overparameterization but rather reflects real heterogeneity in amino acid exchangeabilities over sites.

Finally, the fact that S10C10+G4 trained on the 10,000 sites surpassed taxon-specific models (e.g., Q.yeast+C10+G4) suggests that, when the true model is unknown, and the alignment is sufficiently large, GTRspmix can be a robust alternative. Optimizing the GTRspmix model is computationally expensive, and it may be impractical to perform exhaustive holdout or cross-validation tests as conducted in this study for every dataset. Therefore, for multigene alignments containing dozens or more genes, testing the GTRspmix model with 10 or 30 matrices may offer a practical solution.

### General-purpose GTRspmix Models Outperform Conventional Models

To address cases where model training is challenging due to a limited number of sites (e.g., in single-gene alignments) or computational resource constraints, we estimated general-purpose empirical GTRspmix models (SXXpfamCYY series) using the Pfam database (El-Gebali et al. 2018). Building on the widely used general-purpose CXX profile series (Si Quang et al. 2008), we focused on optimizing GTRspmix models with 10 and 30 exchangeability matrices, based on their good performance with the Eukaryota dataset. The C60 profile set was clustered with C10 and C30 using SPPC clustering. When clustered with C30, the C60 profiles were divided into 29 clusters (one C30 profile had no C60 class assignment and was therefore dropped). Furthermore, one cluster represented by a single low-weight C60 profile (profile 21) was excluded from model estimation, resulting in the final S28pfamC59+G4 model. We also constructed another model by reintroducing the excluded C60 profile 21, which was combined with the POISSON exchangeability matrix, after optimizing the S28pfamC59+G4 model; this model was called S28pfamC60+G4 (see Materials and Methods for a full explanation). The results of clustering are available as Supplementary Figures S2–S4. Each of the C10 and C30 profiles used for SPPC clustering of the C60 profiles was found to be similar to the C60 profiles in the corresponding cluster. Furthermore, we estimated computationally ‘lighter’ models in which each profile of C10, C20, and C30 was assigned a single exchangeability matrix (S10pfamC10+G4, S20pfamC20+G4, and S30pfamC30+G4, respectively). We compared these models and conventional single matrix models using the Pfam test dataset (Table 2).

**Table 2.** Holdout test on Pfam data.

| Model | Train $\ell$ | Train $\Delta\ell$ | AIC | ΔAIC | BIC | ΔBIC | Test $\ell$ | Test $\Delta\ell$ |
| --- | --- | --- | --- | --- | --- | --- | --- | --- |
| Q.pfam+G4 | -52,649,051 | 1,361,306 | 105,817,870 | -2,711,911 | 108,508,122 | -2,656,529 | -53,491,199 | 1,329,396 |
| Q.pfam+C60+G4 | -52,021,157 | 733,412 | 104,562,201 | -1,456,242 | 107,253,064 | -1,401,471 | -52,890,757 | 728,954 |
| G.pfam+C60+G4 | -51,821,049 | 533,304 | 104,162,362 | -1,056,403 | 106,855,182 | -1,003,589 | -52,682,518 | 520,715 |
| S10pfamC10+G4 | -51,753,267 | 465,522 | 104,030,101 | -924,142 | 106,740,011 | -888,418 | -52,620,838 | 459,035 |
| S20pfamC20+G4 | -51,492,016 | 204,271 | 103,511,398 | -405,439 | 106,240,976 | -389,383 | -52,358,730 | 196,927 |
| S30pfamC30+G4 | -51,364,169 | 76,424 | 103,259,504 | -153,545 | 106,008,751 | -157,158 | -52,237,587 | 75,784 |
| S10pfamC60+G4 | -51,381,534 | 93,789 | 103,286,735 | -180,776 | 105,997,163 | -145,570 | -52,242,391 | 80,588 |
| S28pfamC59+G4 | -51,287,745 | — | **103,105,959** | — | **105,851,593** | — | **-52,161,803** | — |
| S28pfamC60+G4<sup>a</sup> | -51,288,360 | 615 | 103,107,190 | -1,231 | 105,852,834 | -1,241 | -52,161,910 | 107 |

<sup>a</sup>S28pfamC60 has one additional component, POISSON+C60pi21, compared with S28pfamC59.
Note: All $\Delta$ values denote differences from the best model.

Consistent with the results based on the Eukaryota dataset, the models with multiple exchangeability matrices outperformed the single-matrix models on the Pfam test dataset as well. Among all models tested, S28pfamC59+G4 achieved the best performance across the three model selection criteria. Notably, even BIC favored S28pfamC59+G4 over the simpler S10pfamC60+G4 model, which differs from the results observed for the Eukaryota dataset (Table S1). This discrepancy may reflect the greater size and diversity of the Pfam dataset, which contains six times more sites than the Eukaryota data (Table 1), potentially exhibiting more complex heterogeneity.

Interestingly, the log-likelihood of S28pfamC60+G4 was slightly lower than that of S28pfamC59+G4, even though the only difference between them is the additional POISSON+C60pi21 class. In principle, if the weight of this extra class was zero, the two models should be equivalent. Indeed, its weight on the Pfam test data was extremely small (1 × 10^−7^), suggesting that the difference in log-likelihood likely arose from minor fluctuations in some parameters such as branch lengths. In subsequent analyses, we used S28pfamC59+G4 and omitted S28pfamC60+G4, although in practical applications, users can compare AIC and/or BIC values between S28pfamC59 and S28pfamC60 and select the better-fitting model.

The improvement in test log-likelihood obtained by moving from Q.pfam+C60+G4 to S28pfamC59+G4 (Δ*ℓ* = 728, 954) was larger than the increase from Q.pfam+G4 to Q.pfam+C60+G4 (Δ*ℓ* = 600, 442). This indicates that introducing the GTRspmix framework yields a gain in model fit greater than that achieved by adding the C60 profile mixture to model PHAS.

Consistent with this observation, S10pfamC10+G4 achieved a higher test log-likelihood than the other single-matrix models with C60 profiles, and S30pfamC30+G4 outperformed S10pfamC60+G4. When computational time or resources are limited, these lighter models (S10pfamC10+G4, S20pfamC20+G4, and S30pfamC30+G4) can serve as practical alternatives. However, S28pfamC59+G4 remained the best-performing model overall, suggesting that, at least for the Pfam dataset, using only 30 profiles is insufficient to capture the full extent of PHAS.

Next, we compared not only the log-likelihoods but also the estimated branch lengths among models using the Pfam test data. We observed a clear trend: the total tree length increased as the profile mixture became more complex, but it decreased as the exchangeability matrix mixture became more complex (Table 3). We note that the upper bound on branch lengths of 10 had a negligible effect on model estimation and evaluation as only a very small number of branches reached this upper limit. Even for the Q.pfam+C60+G4 model, which tended to estimate the longest branches, only 13 branches (0.005%) exceeded 9.9 in length out of 257,536 branches in the Pfam test data. In addition to total tree length, we plotted the estimated branch lengths of all trees and calculated pairwise correlation coefficients between models (Supplementary Fig. S5). Q.pfam+C60+G4 and G.pfam+C60+G4 showed a high correlation in branch length estimates (*r* = 0.9897), although the total tree length under G.pfam+C60+G4 was approximately 84% of that under Q.pfam+C60+G4. S10pfamC60+G4 and S28pfamC59+G4 showed similar total tree lengths and a strong correlation (*r* = 0.9870), indicating that these two models resulted in the most similar branch length estimates among those compared. S20pfamC20+G4 and S30pfamC30+G4 also exhibited high correlations and comparable total tree lengths. The best-fitting model, S28pfamC59+G4, estimated a total branch length that was 79% of that under the Q.pfam+C60+G4 model.

**Table 3.** Branch length inferred on Pfam test data.

| Model | Total Branch Lengths |  |
| --- | --- | --- |
|  | Absolute | Relative (%) |
| Q.pfam+G4 | 59,796 | 69 |
| Q.pfam+C60+G4 | 87,055 | 100 |
| G.pfam+C60+G4 | 73,388 | 84 |
| S10pfamC10+G4 | 60,573 | 70 |
| S20pfamC20+G4 | 64,969 | 75 |
| S30pfamC30+G4 | 67,496 | 78 |
| S10pfamC60+G4 | 69,897 | 80 |
| S28pfamC59+G4 | 68,964 | 79 |

The comparisons of log-likelihoods and branch lengths shown above were performed under a fixed tree topology estimated under Q.pfam+C60+G4. Next, we evaluated how our new models affect tree topology estimation. We conducted tree searches using IQ-TREE v3.0.1 for 100 randomly selected alignments from the Pfam test dataset under each model and compared the resulting ML tree topologies. For each alignment, we calculated the nRF distance between all pairs of models, visualized the distributions across 100 genes using boxplots, and computed the mean values (Supplementary Fig. S6). To assess the stochasticity during tree search, we performed two independent runs with different random seeds under the Q.pfam+C60+G4 model. The average nRF distance between the two runs was 0.0897. This is because for some alignments, the number of sites is low while the number of taxa is very high, causing the IQ-TREE heuristic ML tree estimation algorithm to get caught in local optima. Among the models using C60/C59 profiles, Q.pfam+C60+G4 and G.pfam+C60+G4 showed the most similar topologies (mean nRF = 0.1767), followed by S10pfamC60+G4 and S28pfamC59+G4 (mean nRF = 0.1986). In addition, S20pfamC20+G4 and S30pfamC30+G4 also showed a relatively small distance (mean nRF = 0.1898). In contrast, the largest distance was observed between Q.pfam+C60+G4 and S10pfamC10+G4 (mean nRF = 0.2658).

To investigate which methods tended to estimate more or less similar topologies, we constructed hierarchical clusterings based on the median and mean nRF distance matrices. Because clusterings from the mean and median were identical, we present the median-based UPGMA estimates (Supplementary Fig. S7). The basal clusters completely separate multiple-exchangeability-matrix models from single exchangeability-matrix models. Within these respective groups, models generally clustered according to their parameter complexity, with the simplest model (Q.pfam+G4) forming a singleton cluster. As might be expected, two independent runs of the same model correctly formed a cluster with the highest split frequency over proteins (81%). However, the split frequencies of all other clusters were generally low. This indicates that topological distances between models can be expected to vary substantially over alignments.

#### GTRspmix Models Avoid LBA Artifacts in Three Empirical Analyses

To further evaluate the effect of the newly estimated general-purpose empirical models, we tested them on three empirical concatenated multi-protein datasets that are known to succumb to LBA artifacts under simple single-profile models (Wang et al. 2018; Baños et al. 2024). For each dataset, we were able to compare the log-likelihoods of the likely true and LBA trees between models (Table 4).

**Table 4.** Log-likelihood comparisons between true and LBA trees for empirical concatenated datasets.

| Model | Microsporidia |  |  | Nematode |  |  | Platyhelminthes |  |  |
| --- | --- | --- | --- | --- | --- | --- | --- | --- | --- |
| | True $\ell$ | LBA $\ell$ | $\Delta\ell$ | True $\ell$ | LBA $\ell$ | $\Delta\ell$ | True $\ell$ | LBA $\ell$ | $\Delta\ell$ |
| Q.pfam+G4 | -733,761 | -733,543 | -219* | -728,708 | -728,686 | -22 | -639,941 | -639,850 | -91* |
| Q.pfam+C60+G4 | -716,213 | -716,231 | 18 | -712,609 | -712,640 | 31 | -626,786 | -626,788 | 1 |
| G.pfam+C60+G4 | -713,886 | -713,905 | 20 | -710,751 | -710,799 | 48 | -625,217 | -625,229 | 12 |
| S10pfamC10+G4 | -718,883 | -718,825 | -58* | -715,584 | -715,671 | 87* | -629,573 | -629,608 | 35 |
| S20pfamC20+G4 | -714,188 | -714,187 | -1 | -711,336 | -711,435 | 99* | -625,941 | -625,984 | 43* |
| S30pfamC30+G4 | -713,039 | -713,053 | 13 | -710,094 | -710,195 | 101* | -624,899 | -624,944 | 45* |
| S10pfamC60+G4 | -711,293 | -711,309 | 16 | -708,785 | -708,869 | 84* | -623,800 | -623,840 | 40* |
| S28pfamC59+G4 | -710,892 | -710,900 | 8 | -708,258 | -708,360 | 103* | -623,365 | -623,416 | 51* |

\*Significant difference from Q.pfam+C60+G4 model (Z-test, $p < 0.00238$ after Bonferroni correction).

All models incorporating C60 or C59 profiles favored the true tree over the LBA tree in all three datasets. In contrast, the single-profile model, Q.pfam+G4, consistently supported the LBA tree for all three datasets. The simpler multiple exchangeability/multiple profile models, S10pfamC10+G4, S20pfamC20+G4, and S30pfamC30+G4, all favored the true tree for the Nematode and Platyhelminthes datasets. However, S10pfamC10+G4, S20pfamC20+G4 (but not S30pfamC30+G4) supported the LBA tree for Microsporidia. The log-likelihoods of these empirical data under the true trees differed from those observed in the Pfam test database. In the Pfam analysis, S10pfamC10+G4 outperformed Q.pfam+C60+G4, and S30pfamC30+G4 performed better than S10pfamC60+G4. However, on the empirical LBA datasets, these relationships were reversed, with models incorporating C60 or C59 profiles performing relatively better. This implies that the degree of PHAS in the these datasets is greater than that in the Pfam database.

We performed Z-tests using site-wise log-likelihood differences between the true and LBA trees to assess the statistical significance of the difference in support between the Q.pfam+C60+G4 model and each of the other models (see * in Table 4). Because 21 pairwise tests were conducted, we applied the Bonferroni correction with an adjusted significance level of 0.00238. In the Microsporidia dataset, no significant difference was detected between Q.pfam+C60+G4 and the other models with C60/C59 profiles, suggesting that EHAS had little impact. In contrast, for Nematode and Platyhelminthes, models incorporating multiple exchangeability matrices had significantly different log-likelihood differences compared to the Q.pfam+C60+G4 model, indicating that modeling EHAS contributed to resolving the LBA artifact. Overall, these results suggest that the degree to which failing to model EHAS contributes to LBA artifacts varies across datasets.

To assess model adequacy, we compared the total log-probabilities of the observed sequences under each model for each of the true trees using the ESR-based approach (Table 5). Among all models tested, the highest log-probability of extant (true) sequences was obtained with the S28pfamC59+G4 model, followed by S10pfamC60+G4 and then S30pfamC30+G4, which also outperformed the G.pfam+C60+G4 model. Notably, the ranking of models based on ESR log-probabilities was identical to that based on the log-likelihoods of the true trees. These results indicate that the general-purpose multiple exchangeability profile mixture models were confirmed to perform better from two different aspects.

**Table 5.**
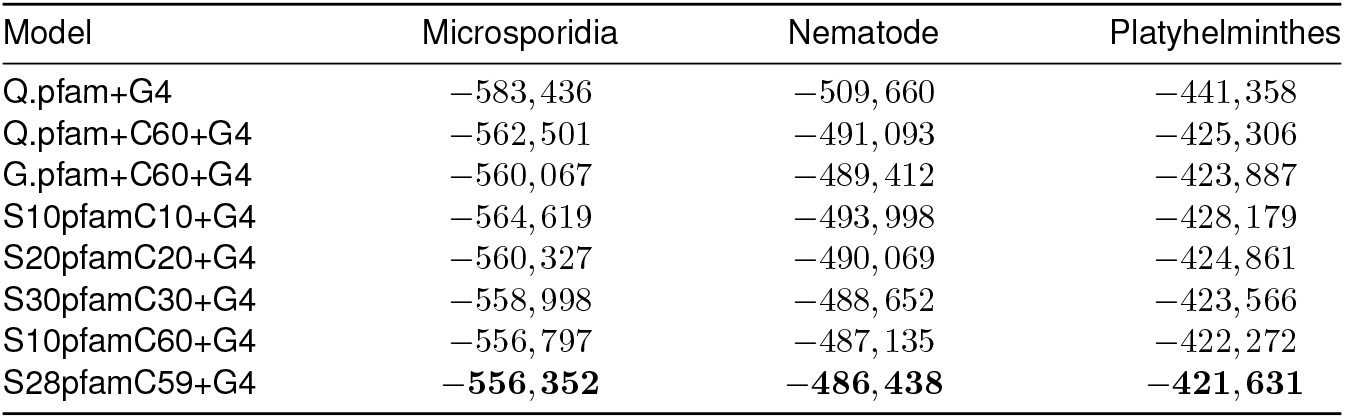
Log-probabilities of the observed sequences in ESR analyses.

We compared the computational time required to fit these newly estimated general-purpose models using the empirical concatenated datasets. For all models, we measured the seconds per iteration in IQ-TREE. All analyses were performed on an AMD Ryzen 7 7700X 8-Core Processor, with three independent runs, and the averages are shown in Table 6. Although S10pfamC60+G4 and S28pfamC59+G4 use multiple exchangeability matrices, the final number of mixture components is the same as the single matrix C60 models (e.g., LG+C60+G4). As a result, their computational costs are comparable to those of conventional C60 models. In contrast, the less complex models—S10pfamC10+G4, S20pfamC20+G4, and S30pfamC30+G4 that were designed for faster analysis—have fewer classes than C60 and accordingly require substantially less computational time, as expected.

**Table 6.** Run time (wall-clock) per iteration on empirical concatenated datasets (sec)

| Model | Microsporidia | Nematode | Platyhelminthes |
| --- | --- | --- | --- |
| Q.pfam+G4 | 0.67 | 0.95 | 0.79 |
| Q.pfam+C60+G4 | 42.58 | 47.83 | 39.89 |
| G.pfam+C60+G4 | 33.03 | 49.83 | 42.61 |
| S10pfamC10+G4 | 8.90 | 7.36 | 9.98 |
| S20pfamC20+G4 | 13.80 | 15.63 | 17.03 |
| S30pfamC30+G4 | 16.79 | 20.42 | 16.88 |
| S10pfamC60+G4 | 38.27 | 41.83 | 35.68 |
| S28pfamC59+G4 | 38.22 | 50.45 | 42.92 |

**Table 7.** Fitness-derived profile and exchangeability matrix distances.

|  |  | Exchangeability matrix distance |  |  |
| --- | --- | --- | --- | --- |
|  |  | low | mid | high |
| Profile distance | low | 0.28 | 0.06 | 0.00 |
|  | mid | 0.06 | 0.23 | 0.05 |
|  | high | 0.00 | 0.05 | 0.28 |

## Discussion

The significant improvements in log-likelihood scores indicates that GTRspmix captures the evolutionary process of proteins more accurately than conventional models (Table S1). Traditionally, profile mixture models like C60 assumed a single global exchangeability matrix across all sites while accounting for site-specific amino acid compositions (Si Quang et al. 2008; Baños et al. 2024). However, our experiments support the presence of EHAS and suggest that a single matrix is insufficient to describe the complex dynamics of protein evolution. Our approach to cluster similar profiles into groups that each share a single exchangeability matrix was motivated by the MutSel model of protein evo-lution, which posits that site-specific substitution rates are determined by the rate matrix under no selection and site-specific relative fitness (see Appendix). Since functional and structural constraints vary by site, fitness landscapes and the resulting exchangeabilities must vary too. Our MutSel simulations confirmed that similar profiles are linked to similar exchangeability matrices as demonstrated by a positive correlation between profile and exchangeability distances (Table 7 and Supplementary Fig. S8).

The effectiveness of our clustering strategy was also compared to a model based on an “anti-clustering” approach, where dissimilar profiles are clustered into groups that share exchangeabilities (see Appendix). The significant drop in likelihood observed from using the anti-clustering-based model clearly demonstrates that grouping similar profiles is more consistent with the underlying evolutionary process than grouping dissimilar ones. These findings not only justify our specific clustering algorithm but also reinforce the fundamental principle that amino acid preferences at sites and amino acid interchange rates are coupled in protein evolution.

Our general-purpose empirical models (SXXpfamCYY series) show substantial improvement in model fit over profile mixture models. In our evaluations using the Pfam database, the log-likelihood improvement achieved by adding GTRspmix to a traditional C60-based model was comparable in scale to the improvement gained by moving from a single-profile model to C60 (Table 2). This indicates that accounting for EHAS is as critical as accounting for site-specific amino acid compositions. Beyond statistical fit, the SXXpfamCYY models influence the estimated branch lengths and tree topologies, effectively mitigating LBA artifacts in datasets where EHAS is particularly pronounced (Table 4 and Supplementary Figs. S5–7). Furthermore, evaluating these general models with ESR revealed its strong potential as a novel model selection criterion (Table 5). For the LBA dataset, the ranking of true sequence log-probabilities under ESR aligned exactly with standard likelihood ranking, and the best-fitting model identified by ESR perfectly matched the one selected by the holdout test on the Pfam data (S28pfamC59+G4). Given that ESR can be applied to any substitution model and dataset, it represents a promising alternative framework for model selection in future studies.

Notably, because the number of rate matrices used in the final mixture remains at 240 (when combined with G4 categories), computational costs are the same as conventional profile mixture models (Table 6). This means that the GTRspmix approach can be easily integrated with other complex mixture models, such as GHOST or MAST (Crotty et al. 2020; Wong et al. 2024). Consequently, these C60-based GTRspmix models represent a practical and broadly applicable alternative to standard C60 analyses for phylogenetic studies of amino acid alignments.

While these general-purpose empirical models provide immediate benefits for standard analyses, the utility of our framework extends further for multi-protein analyses. Users can utilize the GTRspmix script to optimize models specifically tailored to their data. When combined with profile estimation methods like MaM-MAL/MEOW (Susko et al. 2018; Williamson et al. 2025), this allows for a fully data-driven parameterization where both profiles and exchangeability matrices are optimized for the target alignment. Given its broad applicability to any mixture model parameter, our scalable EM approximation approach via soft-partitioning will be essential for making more complex evolutionary models computationally feasible in the future.

Nevertheless, the computational costs associated with model optimization remain a substantial challenge. The execution time of optimizing a GTRspmix model is highly dependent on the number of iterations required for convergence. For instance, fitting an S10M60+G4 model on the Eukaryota training dataset (78 taxa, 38,420 sites) required 11 iterations, taking approximately 5 days on an AMD EPYC 7702 processor using 50 cores. Due to these requirements, simultaneous model optimization and tree searching are currently impractical, and a fixed guide tree is necessary. Given that the guide tree can influence model estimation, it should be selected carefully, perhaps by collapsing poorly supported branches to minimize bias. These computational bottlenecks could be mitigated by implementing GTRspmix natively within IQ-TREE using a true EM algorithm or by adopting rapid estimation methods like PhyloGrad (Lieser et al. 2026).

Looking forward, several extensions could further enhance the framework’s utility. For instance, a fast approximation method similar to the Posterior Mean Site Frequency (PMSF; Wang et al. 2018), which we refer to as Posterior Mean Site Q-matrices, or “PMSQ”, could be developed. By calculating site-specific rate matrices (*Q*) through the posterior-probability-weighted averaging of exchangeability matrices, profiles, and relative rates, PMSQ would enable rapid bootstrap analysis while accounting for EHAS as well as PHAS and RHAS.

Ultimately, the clustering strategy in GTRspmix serves as a bridge between the theoretical fitness model and the practical profile mixture model, enabling the model to capture complex site-specific processes with a manageable computational cost and number of parameters. In conclusion, by accounting for EHAS—a factor overlooked in previous profile mixture models—GTRspmix provides a more realistic representation of protein evolution and offers a robust foundation for estimating reliable phylogenetic trees.

## Acknowledgements

This work was supported by the Natural Sciences and Engineering Research Council of Canada Discovery Grants awarded to A.J.R. (RGPIN-2022-05430) and E.S. (RGPIN-2025-04509); the Moore-Simons Project Call on the Origin of the Eukaryotic Cell, Simons Foundation Grant 735923LPI awarded to A.J.R., E.S., and B.Q.M. (https://doi.org/10.46714/735923LPI); the U.S. National Science Foundation (DEB-2333243) awarded to B.Q.M.; the Na-tional Institutes of Health grants (R01GM096053 and R01GM132499) awarded to D.L.T.; and the Japan Society for the Promotion of Science grants (26KJ0177 and Overseas Research Fellowships) awarded to R.H. Computations in this study were partially performed on the NIG supercomputer at ROIS National Institute of Genetics. The manuscript file uploaded to bioRxiv was generated using the LaTeX template adapted by Stephen Royle available at https://github.com/quantixed/manuscript-templates.

## Data Availability

Data available from Dryad Digital Repository: https://doi.org/10.5061/dryad.7pvmcvf82. Newly estimated general-purpose models (SXXpfamCYY series) are implemented in IQ-TREE version 3.1.3: https://github.com/iqtree/iqtree3. GTRspmix script is available from GitHub repository: https://github.com/HRD-Ryo/GTRspmix.

## Appendix

### Derivation of the MutSel Model

To evaluate the hypothesis that sites belonging to similar profiles would evolve under similar exchangeability matrices, we generated 60 profile-exchangeability matrix pairs under a mutation-selection (MutSel) model and examined whether distances between profiles were associated with distances between their corresponding exchangeability matrices. According to population genetics theory (Kimura 1962; Halpern and Bruno 1998; Yang and Nielsen 2008; Rodrigue et al. 2010; Wang et al. 2014), the relative population-scaled selection coefficient is defined as *F*_*j*_− *F*_*i*_ = 2*Ns*_*ij*_, where *N* is the effective population size, *s*_*ij*_ is the selection coefficient and *F*_*i*_ is the relative fitness of amino acid *i*. Motivated by nearly neutral theory (Ohta 1973), we consider a weak-selection regime in which the population-scaled selection coefficients are typically less than 1 in magnitude. Accordingly, we generated relative fitnesses, *F*_*i*_, from a uniform distribution with a range between 0 to

1. Then, we calculated exchangeabilities and profiles from the relative fitnesses based on the MutSel model. Profiles are calculated by

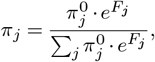

where 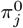is a stationary frequency without selection. In this study, we used an equivalent frequency, 0.05, as 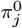, because 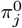 is shared between classes. Exchange-abilities are calculated by

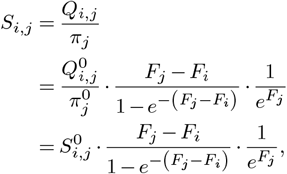

where 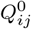and 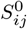 are substitution rate and exchange-ability between amino acids *i* and *j* without selection pressure. We used the POISSON exchangeability matrix as an *S*^0^. And matrix *S* was normalized so that the expected total substitutions would be 1:

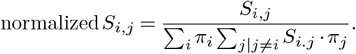

Using generated profiles and exchangeability matrices, we compared profile distances and exchangeability matrix distances. We calculated a 60-by-60 distance matrix by computing the pairwise Euclidean distances between all generated profiles. Similarly, we calculated a 60-by-60 distance matrix of exchangeability matrices. Then, all pairwise distances were shown as a scatter plot (Supplementary Fig. S8) and partitioned into three categories (low, mid, and high) based on tertiles (Table 7).

### Evaluate the Effectiveness of Clustering using Anticlustering

To evaluate the effectiveness of our clustering strategy, which groups similar profiles together, we implemented an “anti-clustering” model, grouping dissimilar profiles. This analysis was conducted using the Eukaryota dataset and the MEOW60 profiles. The anticlustering was performed through the following procedure: we initialized clusters by selecting the two most distant profiles among the set. Then, we iteratively assigned the remaining profiles to groups such that the average distance between the new profile and the existing profiles of the assigned group was maximized. By maximizing Euclidean distance inside profile groups, this approach represents a negative control to verify whether clustering similar profiles is contributing to the model performance.

When evaluated on the Eukaryota test alignment, the anti-clustering-based S10M60+G4 model yielded a log-likelihood of -2,562,650. This performance was significantly inferior to the models based on our proposed clustering strategies: it trailed the SPPC-based S10M60+G4 model (−2,560,390) by 2,260 log-likelihood units (*p* = 1.82 × 10^−27^, paired Z-test).

However, the anti-clustering-based model still performed significantly better than the GTRpmix model (−2,565,525), which has a single exchangeability matrix, with a lead of 2,875 units (*p* = 1.50× 10^−61^, paired Z-test). This result indicates that even an antithetical profile grouping captures exchangeability heterogeneity across sites to some extent. Nevertheless, the significant gap between the similarity-clustering and anti-clustering approaches confirms our conclusion: grouping similar profiles is substantially more representative of the underlying evolutionary process than grouping dissimilar ones.

