## Supplementary Information for "GTRspmix: Capturing Heterogeneity of Exchangeabilities Across Sites to Improve Protein Phylogenetics"

Ryo Harada[1,2,3], Edward Susko[1,4], Thomas K.F. Wong[5,6], Hector Baños[7], Nhan Ly-Trong[5], Robert Lanfear[8], Douglas L. Theobald[9], Bui Quang Minh[5], and Andrew J. Roger[1,2]

[1] Institute for Comparative Genomics, Dalhousie University, Halifax, NS, B3H 4R2, Canada
[2] Department of Biochemistry and Molecular Biology, Dalhousie University, Halifax, NS, B3H 4R2, Canada
[3] Graduate School of Agriculture, Kyoto University, Kyoto, Kyoto, 606-8224, Japan
[4] Department of Mathematics and Statistics, Dalhousie University, Halifax, NS, B3H 4R2, Canada
[5] School of Computing, College of Systems and Society, Australian National University, Canberra, ACT, 2601, Australia
[6] Mathematical Sciences Institute, Australian National University, Canberra, ACT, 2601, Australia
[7] Department of Mathematics, California State University San Bernardino, San Bernardino, CA, 92407, USA
[8] Research School of Biology, College of Science and Medicine, Australian National University, Canberra, ACT, 2601, Australia
[9] Department of Biochemistry, Brandeis University, Waltham, MA, 02453, USA

June, 2026

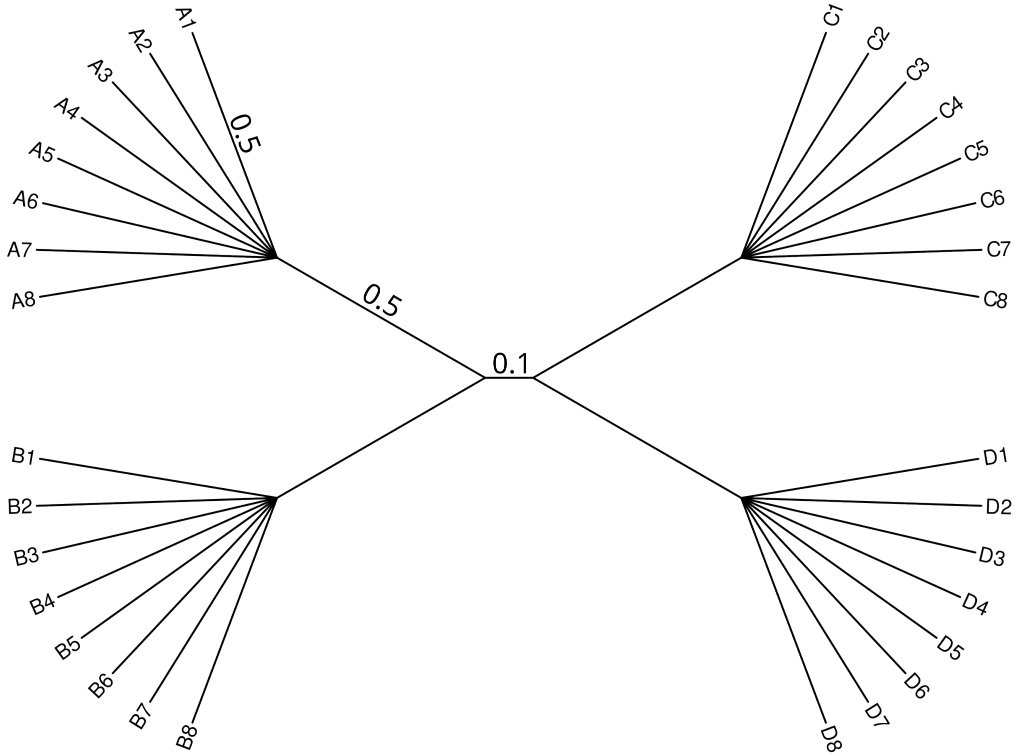


Figure S1: Tree of 32 taxa used to generate sequences in simulations for evaluating overparameterization. The central branch has a length of 0.1, while all other branches have a length of 0.5. The corresponding tree in Newick format is as follows:

```
(((A1:0.5,A2:0.5,A3:0.5,A4:0.5,A5:0.5,A6:0.5,A7:0.5,A8:0.5):0.5,
(B1:0.5,B2:0.5,B3:0.5,B4:0.5,B5:0.5,B6:0.5,B7:0.5,B8:0.5):0.5):0.1,
(C1:0.5,C2:0.5,C3:0.5,C4:0.5,C5:0.5,C6:0.5,C7:0.5,C8:0.5):0.5,
(D1:0.5,D2:0.5,D3:0.5,D4:0.5,D5:0.5,D6:0.5,D7:0.5,D8:0.5):0.5);
```

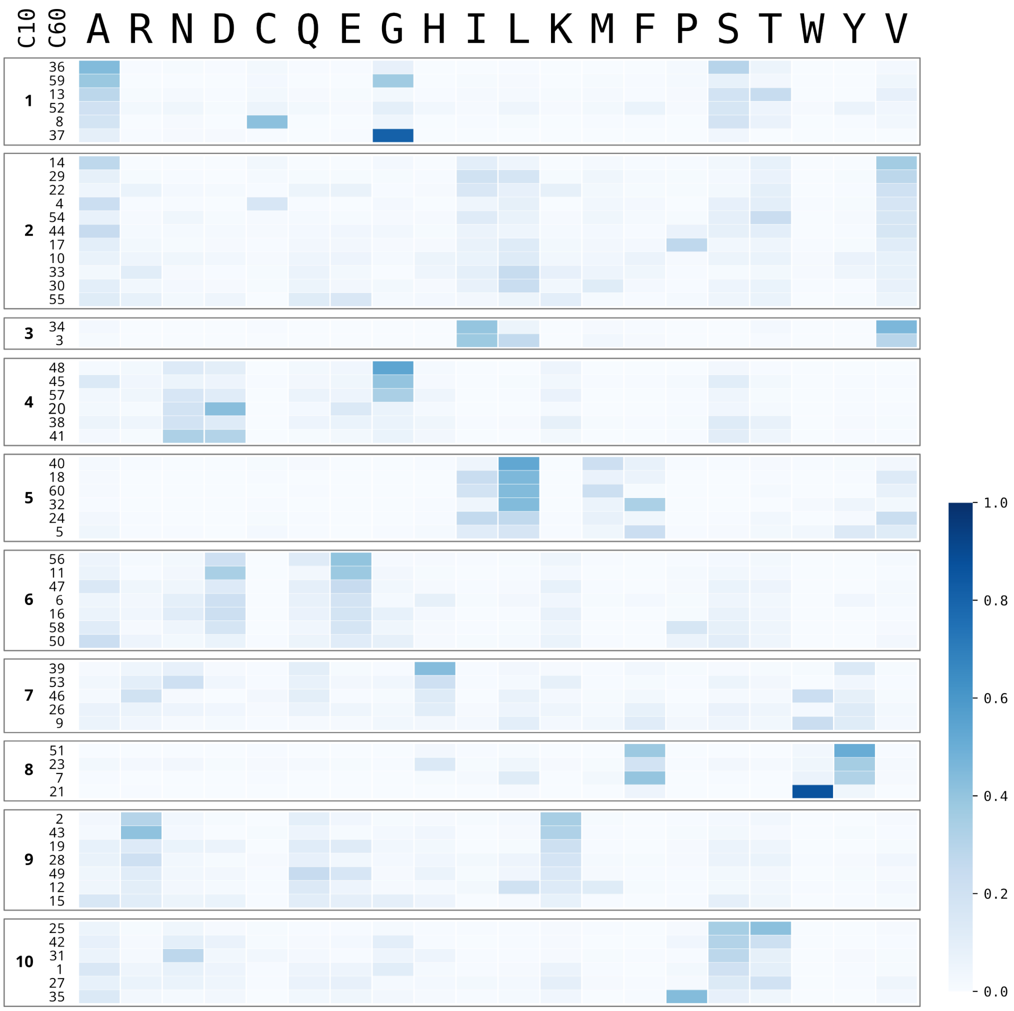


Figure S2: Results of the SPPC clustering used in the S10pfamC60 model. The 60 profiles included in C60 are shown as a heatmap. The C60 profiles are grouped into 10 clusters, and the left side of the figure indicates the corresponding C10 profile indices and C60 profile indices. The right bottom color scale is showing the frequency of amino acids.

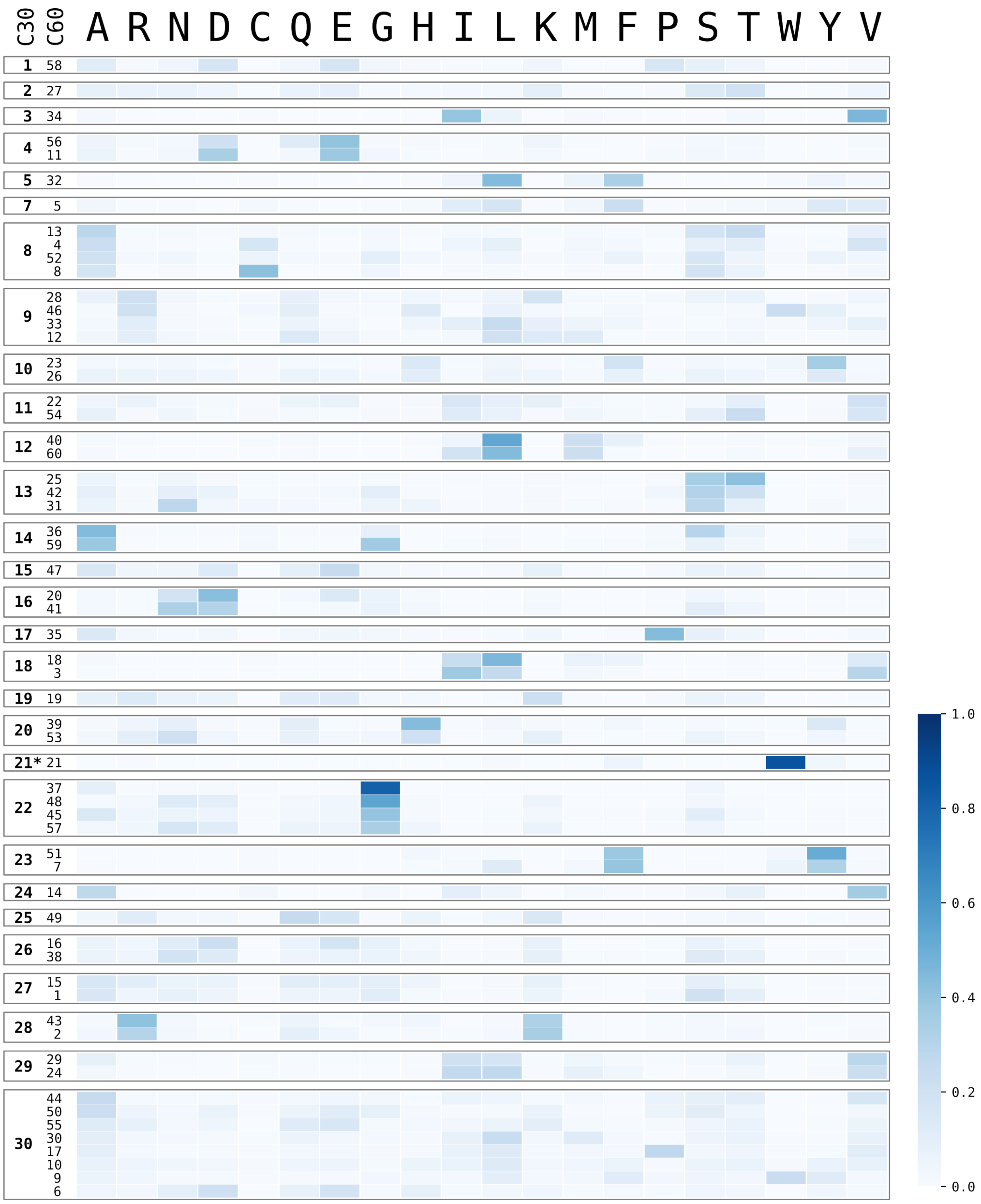


Figure S3: Results of the SPPC clustering used in the S28pfamC59 model and the S28pfamC60 model. The 60 profiles included in C60 are shown as a heatmap. The C60 profiles are grouped into 29 clusters, and the left side of the figure indicates the corresponding C30 profile indices and C60 profile indices. The right bottom color scale is showing the frequency of amino acids. *C60pi21 was removed from model optimization and not used in S28pfamC59 model.

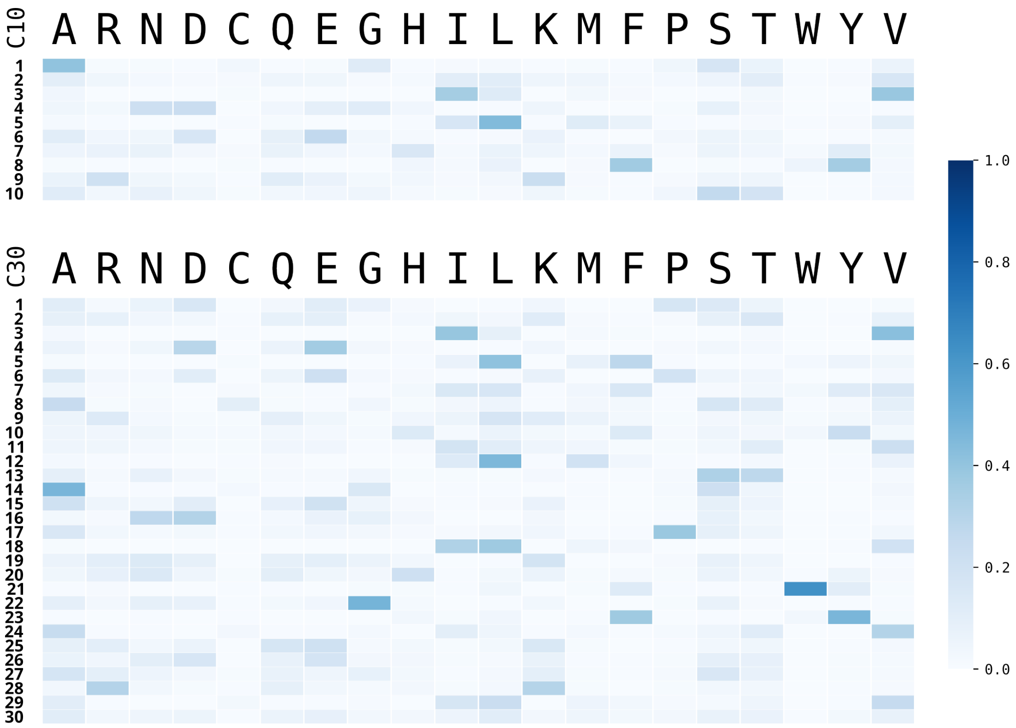


Figure S4: Profiles in C10 and C30. The right bottom color scale is showing the frequency of amino acids.

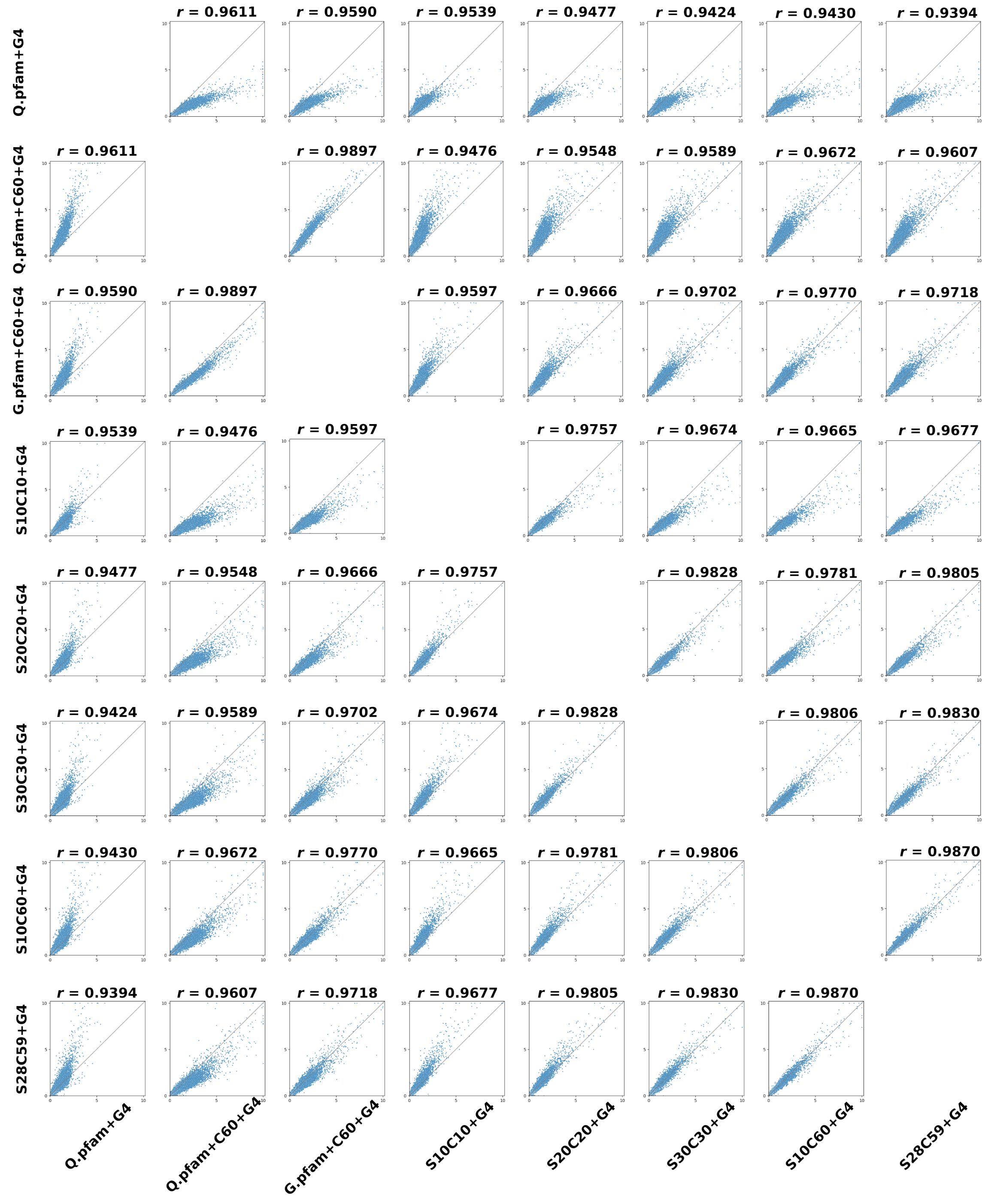


Figure S5: Comparison of branch lengths between all pairs of models. Each scatter plot shows branch lengths from the model on the x-axis versus the model on the y-axis. The correlation coefficient ($r$) is indicated above each plot.

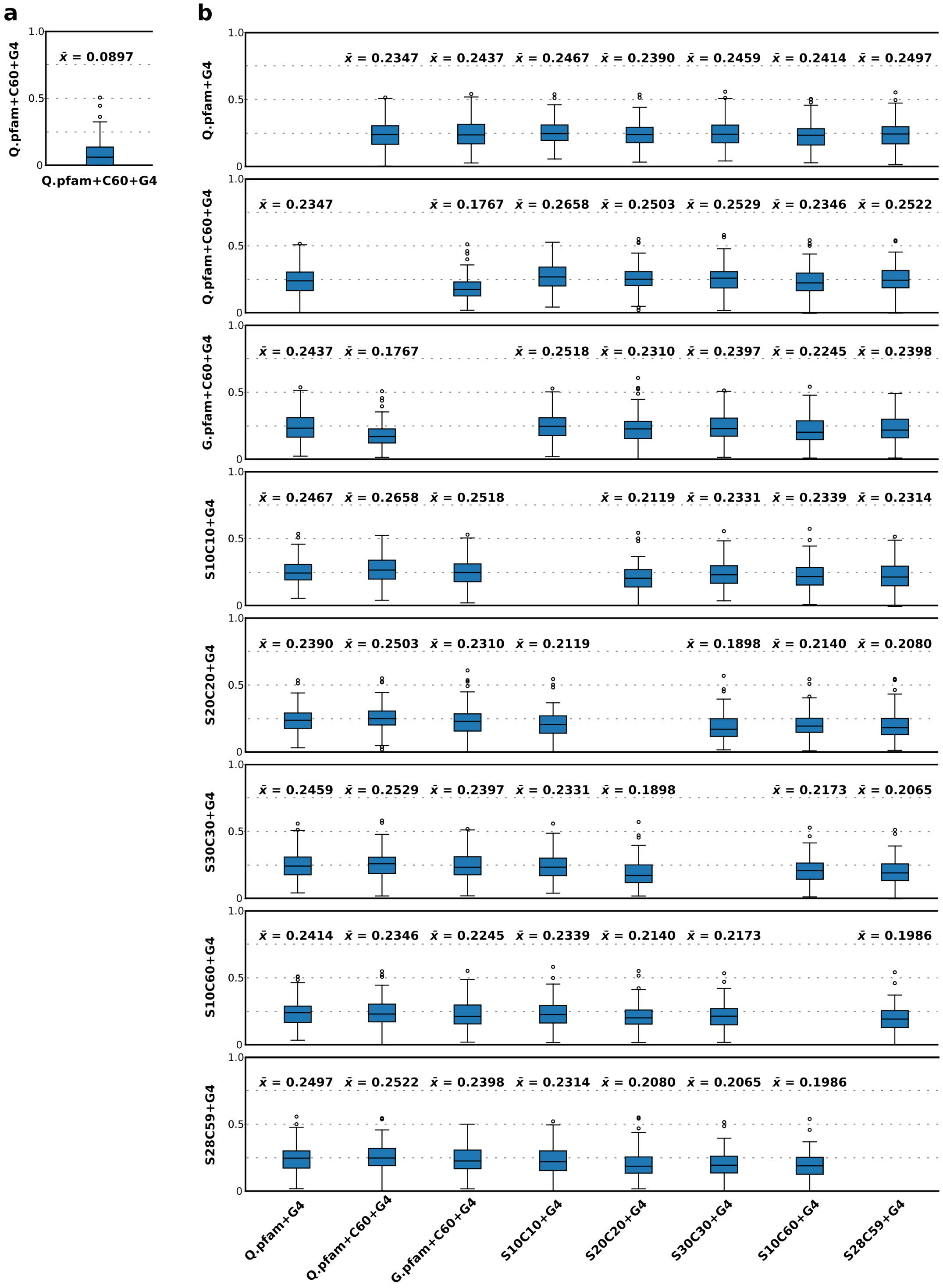


Figure S6: Comparison of normalized Robinson-Foulds (nRF) distances between all pairs of models. a) Comparison of two independent tree searches under the Q.pfam+C60+G4 model. b) Each boxplot shows the distribution of nRF distances across 100 genes for the model on the x-axis versus the model on the y-axis. The average nRF distance is indicated above each plot.

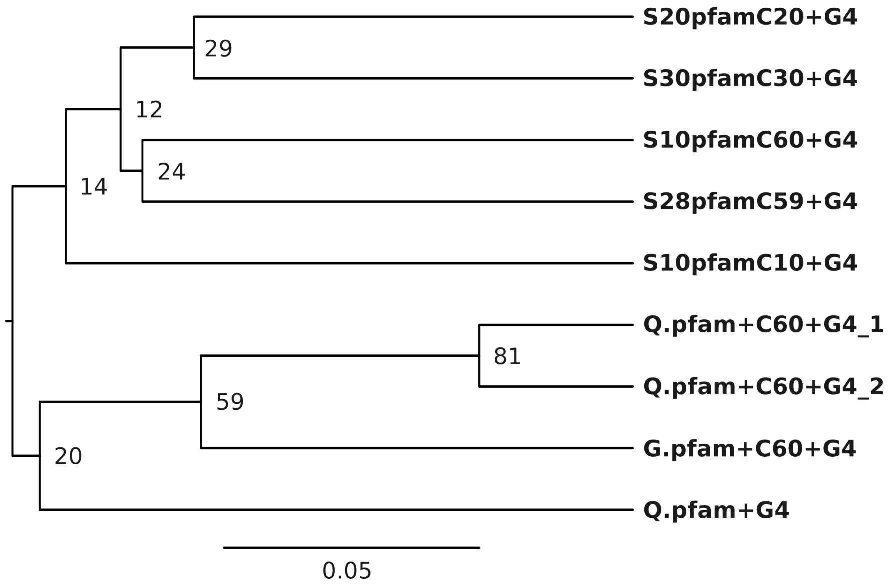


Figure S7: Hierarchical clustering (Unweighted Pair Group Method with Arithmetic Mean; UPGMA) of the models based on the element-wise median normalized Robinson-Foulds (nRF) distance matrix derived from 100 alignments. Numbers at branches indicate the split frequencies (%) calculated from the 100 individual gene trees. The scale bar represents the median nRF distance.

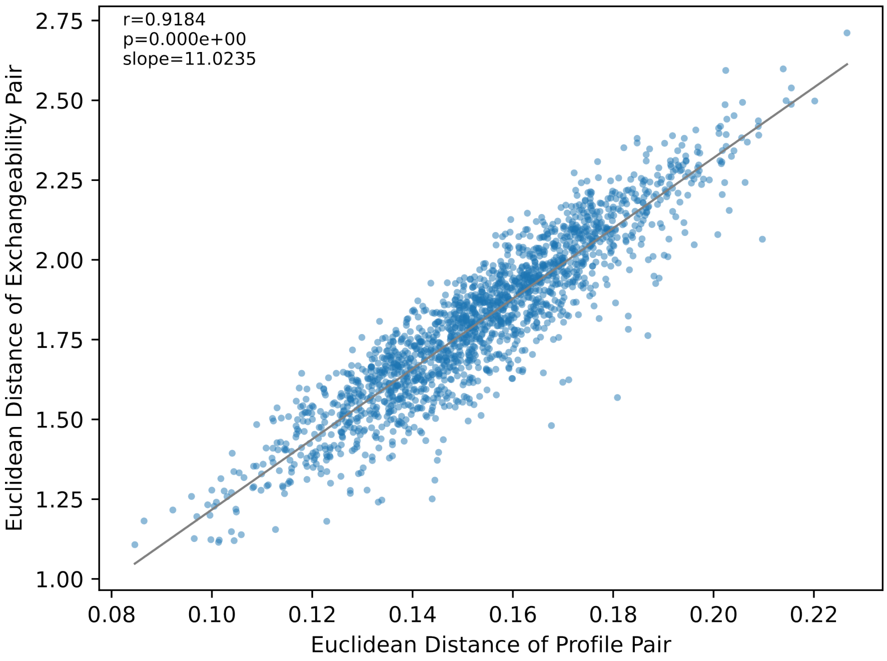


Figure S8: Relationship between profile distances and exchangeability matrix distances. The x-axis and y-axis are showing the Euclidean distance of profile pair and distance of exchangeability matrix pair, respectively. The diagonal line represents the fitted linear regression, and the upper-left inset reports Pearson's correlation coefficient, associated p-value, and regression slope.

Table S1: Holdout test on Eukaryota data

| Model | Clustering | Matrices | Train $\ell$ | AIC | BIC | Test $\ell$ |
|---|---|---|---|---|---|---|
| POISSON+M60+G4 | NA | 0 | $-2,588,052$ | $5,176,530$ | $5,178,352$ | $-2,594,346$* |
| GTRpmix+M60+G4 | NA | 1 | $-2,558,330$ | $5,117,464$ | $5,120,904$ | $-2,565,525$* |
| S02M60+G4 | SPPC | 2 | $-2,555,593$ | $5,112,368$ | $5,117,425$ | $-2,563,363$* |
| S02M60+G4 | Kmeans | 2 | $-2,555,456$ | $5,112,094$ | $5,117,151$ | $-2,563,232$* |
| S04M60+G4 | SPPC | 4 | $-2,553,277$ | $5,108,492$ | $\mathbf{5,116,783}$ | $-2,561,760$* |
| S04M60+G4 | Kmeans | 4 | $-2,553,677$ | $5,109,292$ | $5,117,583$ | $-2,562,256$* |
| S06M60+G4 | SPPC | 6 | $-2,552,622$ | $5,107,938$ | $5,119,463$ | $-2,561,697$* |
| S06M60+G4 | Kmeans | 6 | $-2,551,344$ | $5,105,382$ | $5,116,907$ | $-2,560,570$ |
| S08M60+G4 | SPPC | 8 | $-2,550,738$ | $5,104,926$ | $5,119,686$ | $-2,561,159$* |
| S08M60+G4 | Kmeans | 8 | $-2,550,714$ | $5,104,878$ | $5,119,638$ | $-2,561,370$* |
| S10M60+G4 | SPPC | 10 | $-2,549,462$ | $5,103,130$ | $5,121,124$ | $-2,560,390$ |
| S10M60+G4 | Kmeans | 10 | $-2,550,173$ | $5,104,552$ | $5,122,546$ | $-2,561,333$* |
| S20M60+G4 | SPPC | 20 | $-2,545,357$ | $5,098,700$ | $5,132,865$ | $-2,560,453$ |
| S20M60+G4 | Kmeans | 20 | $-2,546,081$ | $5,100,148$ | $5,134,313$ | $-2,560,655$ |
| S30M60+G4 | SPPC | 30 | $-2,543,029$ | $5,097,824$ | $5,148,161$ | $\mathbf{-2,560,142}$ |
| S30M60+G4 | Kmeans | 30 | $-2,542,899$ | $5,097,564$ | $5,147,901$ | $-2,560,243$ |
| S37M60+G4 | SPPC | 37 | $-2,540,991$ | $5,096,394$ | $5,158,051$ | $-2,560,454$ |
| S40M60+G4 | Kmeans | 40 | $-2,540,848$ | $5,097,242$ | $5,163,750$ | $-2,560,424$ |
| S47M60+G4 | SPPC | 47 | $-2,537,919$ | $5,094,030$ | $5,171,858$ | $-2,561,026$* |
| S50M60+G4 | Kmeans | 50 | $-2,538,106$ | $5,095,538$ | $5,178,218$ | $-2,560,528$ |
| S60M60+G4 | NA | 60 | $-2,535,407$ | $\mathbf{5,093,920}$ | $5,192,771$ | $-2,561,037$* |

*Significant difference from S30M60+G4 model with SPPC (Z-test, $p < 0.0025$ after Bonferroni correction).
